# Investigating the impact of carbamazepine on tomato plant metabolism using genome-scale metabolic modelling

**DOI:** 10.1101/2025.02.03.635865

**Authors:** Samyuktha Srinivasan, Karthik Raman, Smita Srivastava

## Abstract

A comprehensive mechanistic analysis of emerging pharmaceutical pollutants’ stress response in plants is needed to understand its chronic impact on food-chain contamination and agricultural productivity. To unravel this at systems-level, the current study employs insights from green-liver concept and establishes the utility of constraint-based modelling approach for elucidating perturbations in a plant’s metabolism due to pharmaceutical stress.

In this study, the stress response of an emerging recalcitrant anticonvulsant pollutant, carbamazepine (CBZ), was simulated in tomato crop under phototrophic conditions. For this, an updated genome-scale metabolic model of tomato leaf (CBZ_*i*SL3433) was developed and augmented with CBZ transformation reactions based on the green-liver concept. The model was able to capture energy and co-factor competition-induced biomass reduction in presence of CBZ stress. Further, the study provides an *in silico* mechanistic proof for abiotic stress response induced by CBZ in tomato with altered flux states in nutrient assimilation, synthesis of key precursors of leaf biomass and secondary metabolites. Additionally, to extend the applicability of model, potential ameliorative effects of biostimulants such as proline, spermine, glycerol, and ethanol were investigated through model predictions. Through systematic computational analysis, 154 significantly altered reactions were identified in the presence of CBZ stress, of which 92 % of reactions were ameliorated with biostimulants. Amino acid biosynthesis was found to be the most significantly altered pathway under CBZ stress in the presence of biostimulants.

Overall, the proposed framework can aid in screening and developing rational strategies to maintain agricultural yields amid rising plant stress due to such anthropogenic pollutants.

---

1. **Introduction**

In the past three decades, pharmaceutical residues have been identified as emerging contaminants that are pseudo-persistent in the environment owing to their low removal efficiency in conventional water treatment plants (Patel et al. 2019). With approximately 70 % of freshwater required for agriculture and to overcome the global water stress with a projected 53 % increased demand for water by 2050, increased efforts are being made to reuse wastewater for agriculture (Poudel et al. 2023). However, the use of treated wastewater for irrigation exposes agricultural soil and crops to biologically active pharmaceuticals and their bioaccumulation in the food chain. Pharmaceuticals are designed to be biologically active and hence pose the risk of high ecological imbalance and are also known to aggravate antimicrobial resistance (Wang et al. 2019). This impact can be compounded by the projected increase in the usage of reclaimed water worldwide and the global rise in pharmaceutical production and consumption (Nguyen et al. 2023). Therefore, understanding the impact of pharmaceutical pollutants present in reclaimed water used in agriculture can directly or indirectly address sustainable developmental goals (SDGs), majorly SDG 6 (improved wastewater management), and SDG 15 (sustainable agricultural practices development).

Pharmaceutical pollutants have been recognized for their disruptive effects on plant homeostasis (Christou et al. 2018). The extent of impact on plants depends on the soil properties, temporal variations, and physio-chemical properties of the pharmaceutical(s) and their synergistic and antagonistic effects when present as mixtures. Moreover, comprehensive mechanistic insights into both pharmacokinetics and pharmacodynamics in plants are limited. Also, relying only on whole-plant studies for investigating pharmaceutical stress response taking into account all these variations, while informative, is arduous and time-consuming. Thus, there is a need for a more rational framework to flag the type of pharmaceuticals that may present a greater risk of affecting major plant functions in terms of metabolic perturbations (Garduño-Jiménez and Carter, 2024). Such an approach will enable comprehensive scrutiny of the critical metabolic changes induced in plants by such pharmaceutical pollutants for focused experimental investigations. Thus, this will help in reducing trial and error approach, thereby minimizing time, cost, and resource utilization.

For this, we propose a framework for exploiting the predictive power of GEMs (GEnome-scale Metabolic models) approach. The GEM of an organism is an *in silico* stoichiometric model that consists of the complete metabolic inventory of the organism. Though a eukaryotic plant GEM is more complex due to multi-tissue and multi-organ metabolism, they have been proven to be capable of robust predictions of metabolic states observed during abiotic stress response (Williams et al. 2010; Lakshmanan et al. 2016; Herrmann et al. 2019; Wanichthanarak et al. 2020; Chowdhury et al. 2023). Further, with respect to drug metabolism, GEM has been used for understanding drug-induced perturbations on liver toxicity (Cordes et al. 2018) and also for investigation of the effects of diet on drug metabolism in humans (Sahoo et al. 2015). However, xenobiotic stress response in plants has not yet been explored using GEM.

To fill this gap, we propose to adapt ‘green-liver’ concept (Sandermann et al. 1977; Scheel and Sandermann Jr. 1977), recognizing that plant xenobiotic metabolism closely resembles that of animal liver metabolism. Pharmaceuticals are among the class of chemicals with rich data in terms of toxicology in mammalian systems (Malchi et al. 2022). The sequence of xenobiotic metabolism has been reported to be homologous in terms of cDNA sequence, enzyme classes, and metabolite patterns between plant vs animal liver systems with avenues of applications for pharmacovigilance in plants (Coleman et al. 1997; Carter et al. 2019; Malchi et al. 2022; Garduño-Jiménez and Carter, 2024).

In the present study, emerging recalcitrant pollutant carbamazepine (CBZ) has been selected as a representative pharmaceutical for investigation due to its ubiquitous presence and its teratogenic and antimicrobial resistance-inducing nature (Wang et al. 2019). CBZ is known to be easily taken up by various plant species, and its exposure has been known to trigger a typical abiotic stress response in plants (Gorovits et al. 2020). However, a comprehensive systems-level understanding of CBZ-induced metabolic alterations is not yet explored in plants (Mascellani et al. 2023; Wei et al. 2023). For this, the effect of CBZ is proposed to be investigated in one of the majorly consumed crop, tomato (*Solanum lycopersicum*). With the available information on CBZ transformation (Riemenschneider et al. 2017) along with the existence of GEM for tomato (Gerlin et al. 2022), a rational approach for evaluating CBZ-specific stress response could be investigated in this agronomically important crop.

Moreover, the addition of phytoprotectant metabolites as biostimulants has been considered as non-invasive effective way to improve abiotic stress tolerance (Shiade et al. 2024); yet, there are gaps in systems-level understanding of its mechanism of action. Also, their effect on alleviating pharmaceutical stress in plants remains unexplored. Thus, the current study was undertaken to capture pharmacodynamics of CBZ in tomato using genome-scale metabolic modelling and establish it as a predictive tool for rapid screening of potential nutritional supplements that can serve as biostimulators of CBZ stress tolerance.

## 2. Methods

### 2.1 Genome scale metabolic model of tomato leaf: Curation and modification of tomato GEM

The GEM module (Gerlin et al. 2022) of the multi-organ metabolic model of Virtual Young TOmato Plant (VYTOP) was adapted to simulate the CBZ stress response. The distribution of tissue-specific CBZ pharmacokinetics data was found to be limited in literature. Therefore, GEM was preferred to focus on investigations of perturbations in a single cell. Among the vegetative tissues, having the photosynthetic machinery, the leaf is one of the most metabolically active parts with respect to pharmaceutical stress tolerance. With a log *K*_ow_ of 2.45 (Ravichandran and Philip 2022), signifying its neutral state, CBZ-specific transformation reactions have been reported to occur extensively in leaf (Schröder 2007, Riemenschneider et al. 2017). Also, Lu et al. (2024) reported that leaf edges are primary site for CBZ transformation with elevated oxidative stress response in tomato plants exposed to CBZ. Based on this rationale, among the different tissues, the GEM module was converted as a leaf module for pharmacodynamic evaluation. Improvements to the base model connectivity and the input constraints to the model are detailed in Supplementary Text section S1.1. The details of the reactions added and improved model in SBML file format have been provided in supplementary as ‘Supplementary_1.xlsx’ and ‘CBZ_*i*SL3433.xml’ respectively.

### 2.2 Curation of CBZ pharmacokinetics module

The overview of the CBZ pharmacokinetic (CBZ_PK) module developed in the current study is depicted in Figure 1. According to Riemenschneider et al. (2017), in tomato plant, as a part of phase-I metabolism, CBZ has been reported to be transformed via epoxidation to carbamazepine-10,11-epoxide (CBZE) and hydroxylated at different positions to 2-Hydroxycarbamazepine (2-OH-CBZ), 3-Hydroxycarbamazepine (3-OH-CBZ) and 4-Hydroxycarbamazepine (4-OH-CBZ) majorly via cytochrome-P450 mediated transformation. Further, via epoxide hydrolase, CBZE is reported to be converted predominantly to 10,11-Dihydroxycarbamazepine (DiOH-CBZ), which has been known to be converted to oxcarbazepine. Next, with phase-II metabolism, most of the phase-I metabolites (CBZE, 2-OH-CBZ, 3-OH-CBZ, 4-OH-CBZ) have been observed to undergo O-glycosyltransferase mediated conjugation leading to the formation of glucose conjugated CBZ. Also, glutathione-S-transferase (GST) mediated conjugation of glutathione to CBZE has been reported as a part of phase-II metabolism. Further, CBZE conjugates of glutathione, are reported to be transported to vacuole via ATP-dependent pumps, where they have been reported to be degraded to cysteine conjugates by glutamyl transpeptidase as a part of phase-III metabolism. The information on the representation of CBZ transformation reactions is detailed in Supplementary Text section S1.2, and the details for this module are provided in the supplementary Excel file ‘Supplementary_2.xlsx’.

**Figure 1:**
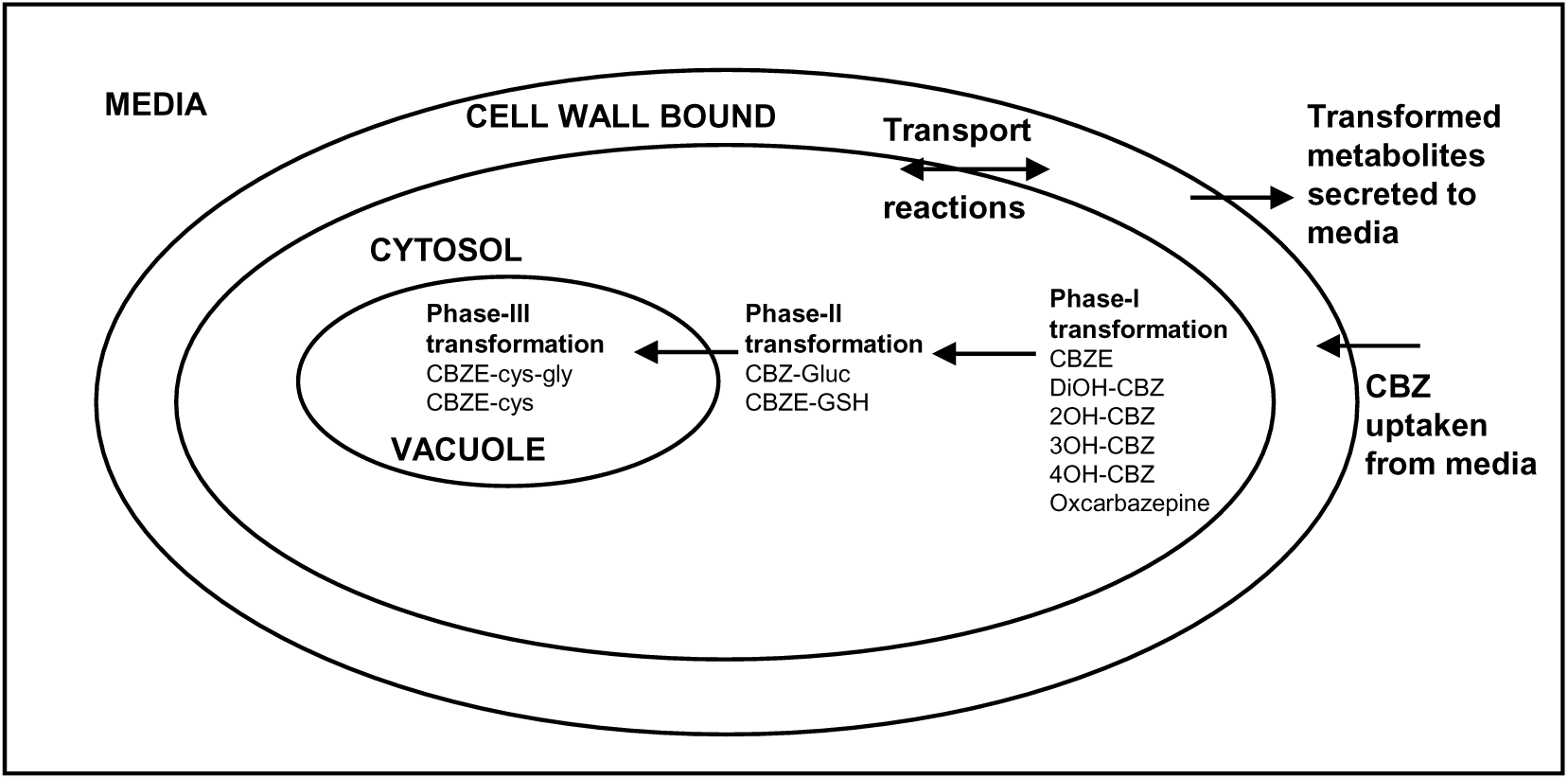
A schematic representation of the CBZ_PK module. Figure illustrates a schematic representation of the various key steps included for CBZ detoxification in the model.

To evaluate the balanced model with the CBZ_PK module, a MEMOTE (Lieven et al. 2020) report was generated and compared to the base model version. Further, the updated model’s reproducibility was evaluated with FROG test suite utilizing the ‘FBC Curation MATLAB Toolbox’ (Raman et al. 2024). The FROG test suite conducts a systematic assessment of the consistency of model simulations across different computational environments, focusing on Flux variability analysis, Reaction deletion, Objective function values, and Gene deletion.

Phototrophic, stress, and mutant conditions were simulated and verified as per the prior version of tomato model (Gerlin et al. 2022). In addition, sanity checks (Heirendt et al. 2019) were implemented to test for the quality of the addition of CBZ_PK module to the updated tomato leaf model.

### 2.3 Simulation for predictions of CBZ pharmacodynamics

#### 2.3.1 Simulation of the effect of CBZ stress on specific growth rate

The effect of CBZ was simulated with a pharmacokinetic module of CBZ along with an exchange reaction for CBZ input and corresponding exchange fluxes simulating the output of CBZ-transformed metabolites. CBZ uptake was enforced by fixing both the upper and lower bounds of the CBZ exchange reaction to study the effect of CBZ stress. The simulations were done under phototrophic mode to represent a physiologically relevant stress condition. Due to the non-availability of information on non-growth associated maintenance (NGAM) and change in biomass precursors under the presence of CBZ, similar values as mentioned in Table S1 for NGAM and the same leaf biomass reaction were used for simulating CBZ stress. With increasing CBZ uptake rate in the range of 0 to 1000 mmol gDW^-1^ h^-1^ under phototrophic conditions, as mentioned in Table S1, the specific growth rate was evaluated using flux balance analysis (FBA) (*l1*-norm) with maximizing biomass reaction as the objective function. The *l1*-norm parameter signifying simultaneous minimization of the sum of flux in the network, the maximum possible specific growth rate with a minimal set of active reactions was determined using FBA.

#### 2.3.2 Simulation for identification of altered metabolism with CBZ stress

Flux variability analysis (FVA) was performed to determine the optimal flux solution space for each reaction with maximizing biomass reaction as the objective function. Based on FVA, parameters such as Jaccard similarity index have been reported as useful metrics for identifying differentially altered reactions (Nanda et al. 2020). In this study, to find differentially altered reactions with CBZ stress, FVA was estimated under two conditions: Control (Uptake flux of CBZ of zero) and CBZ_IC50 (Uptake flux of CBZ inducing 50 % reduction in specific growth rate compared to control conditions). Further, Similarity Index (SI) was estimated based on Jaccard index as per Equation 1 to represent a quantitative measure for similarity between two model conditions. With the minimum and maximum flux ranges obtained for all reactions from FVA for both conditions, SI was computed using COBRA Toolbox function, ‘fvaJaccardIndex’. Reactions with zero SI implying no overlap in flux solution spaces between two conditions was considered to represent significant metabolic alterations.

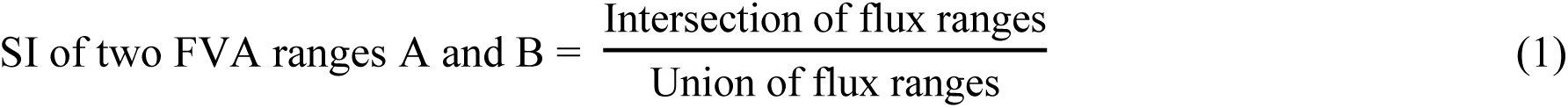

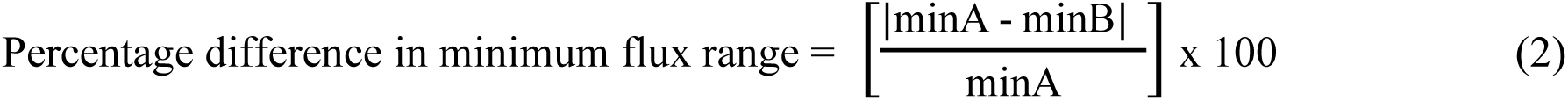

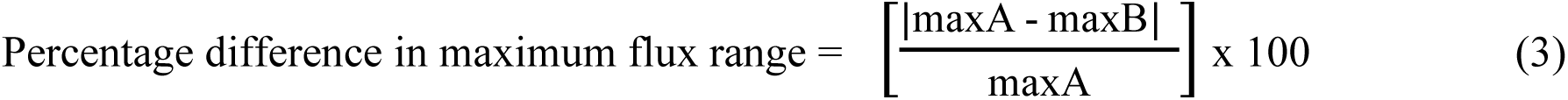

where,

A: (minA, maxA) flux range under Control condition

B: (minB, maxB) flux range under CBZ_IC50 condition Intersection of flux ranges: min(maxA,maxB) − max(minA,minB) Union of flux ranges: max(maxA,maxB) − min(minA,minB)

The reactions were then clustered into pathways on the basis of their metabolic functionalities to investigate the alterations with CBZ stress over the network. Further, for pathways known to be altered under pharmaceutical stress in plants, additionally the percentage difference in flux ranges were calculated as per Equations 2 and 3 and the reactions were categorized as per Figure S2. The details of the reactions and the pathways investigated in this study are provided in supplementary excel file ‘Supplementary_1.xlsx’.

All simulations were performed in MATLAB R2021b (MathWorks Inc., USA) using the COBRA Toolbox v3.0 and Gurobi solver v9.5 as the linear programming solver.

### 2.4 Investigating the impact of biostimulants for improving the CBZ stress tolerance in plants

#### 2.4.1 Simulating the effect of biostimulant on specific growth rate in presence of CBZ stress

Novel sustainable agricultural practices like the application of exogenous biostimulants have been reported to be a promising lead to tackle abiotic stress. Biostimulants include products of organic/inorganic origin and/or microbes known to stimulate plant nutrition process independent of their nutrient content (Jiménez-Arias et al. 2021). Foliar addition of biostimulants has been known to improve nutrient uptake, improve photosynthetic rates, and aid in tolerating ROS thus improving plant growth under abiotic stress like drought, temperature, salinity and heavy metal (García-García et al. 2020). Despite the growing interest in biostimulants, a more comprehensive understanding of their causal and functional mechanisms is still lacking (Van Oosten et al. 2017). Further, in terms of pharmaceutical stress tolerance, the role of biostimulants has not yet been explored.

For this, the ameliorative effect on growth rate of leaf biomass in the presence of CBZ was evaluated by mimicking the exogenous addition of the following four common biostimulants under each category of: an amino acid – proline, a polyamine – spermine, an alcohol – ethanol and a polyol – glycerol. To mimic the exogenous foliar addition, an exchange reaction for the corresponding biostimulant was added and its effect was compared in the presence and absence of CBZ stress. With increasing uptake rates of biostimulants, FBA (*l1*-norm) with maximizing biomass reaction as objective function was estimated under Control and CBZ_IC50 conditions.

#### 2.4.2 Simulation for the ameliorative effect of biostimulants in the presence of CBZ stress

Similar to Section 2.3.2, FVA and SI were estimated to identify significantly altered reactions with biostimulants. As per Figure S2, the altered reactions were categorized for two conditions, CBZ_IC50_Biostimulant and CBZ_IC50 representing effect of the presence and absence of biostimulant under CBZ stress respectively. Further, the effect of biostimulants was evaluated on the production capability of individual biomass components and transformed metabolites of CBZ. The details of the analysis are provided in Supplementary Text section S1.4.

## 3. Results and Discussion

### 3.1 CBZ Pharmacokinetic module based on green-liver hypothesis

In the current study, the developed CBZ_PK module was based on experimental proof of transformed metabolites reported in tomato with phase-I reactions adapted from animal liver. This was based on homologous pharmaceutical metabolism between plants and animal liver as per the green-liver concept. CBZ in animal liver systems is known to be detoxified by Cytochrome-P450 majorly via CYP3A4 mediated phase-I transformation, UDP-Glucuronosyltransferase mediated phase-II transformation, and finally, the converted relatively more hydrophilic version of the detoxified transformation product is excreted from the system (Thorn et al. 2011; Whirl-Carrillo et al. 2021).

In tomato, the equivalent of human CYP3A4 was found using BLAST, Cytochrome-P450 711A1 (BLASTP: Identity: 29 %; Positives: 49 %). Therefore, reactions pertaining to that were added in a similar way as for the animal-liver system. Also, potential variations specific to plant systems were considered for adding phase-II and phase-III metabolism. It has been reported that plant UGTs conjugate a variety of molecules with UDP-Glucose rather than UDP-Glucuronate as substrate (Meech et al. 2012). Thus, human UDP2B7 (UDP-Glucuronosyltransferase) equivalent tomato UDP-glycosyltransferase 76-B1 (BLASTP: Identity:33 %; Positives: 51 %) based detoxification of phase-II transformations of CBZ was added. Further, unlike the excretion of phase-III metabolites as observed in animal system, glutathione conjugated phase-II metabolites were transported to vacuole as observed in plant systems (Coleman et al. 1997).

In support of the green-liver concept, upregulation of genes of Cytochrome-P450, epoxide hydrolases and *O*-glucosyltransferase enzymes have been reported in tomato leaves exposed to CBZ (Lu et al. 2024). In lettuce roots exposed to CBZ, based on a proteomics study, cytochrome P450 81Q32 and cytochrome P450 82C4 proteins have been observed to be significantly increased in comparison to the absence of CBZ (Leitão et al. 2021a). Also, in *Arabidopsis thaliana* transcriptomics study, in roots exposed to emerging contaminants, ibuprofen (Landa et al. 2017), naproxen and praziquantel (Landa et al. 2018), several transcripts of cytochrome-P450, UDP-glycosyltransferases and GST were reported to be upregulated. Further, based on docking studies for mycotoxin aflatoxin B1, it has been reported that *Zea mays* cytochrome-P450 (CYP450 81D11) mirrored its binding to that of human cytochrome-P450 (CYP450 1A2) with sequence identity of 30.65 % (Righetti et al. 2021). Interestingly, the transformation of *Medicago sativa* plants with the human cytochrome-P450 (*CYP2E1*) and GST genes improved phytoremediation of mercury-trichloroethylene contamination (Zhang et al. 2013).

The updated tomato leaf model with CBZ_PK module added based on the green-liver concept consists of 2145 metabolites and 2328 reactions (Figure 2). Based on the MEMOTE report, the updated model with the CBZ_PK module (CBZ_*i*SL3433) has an overall score of 51 % with improved consistency in comparison to base model. Notably, in the updated model, charge balance was improved from 55.4 % to 97.1 % and mass balance was improved from 81.8 % to 95.5 % in comparison to the base model. The results for the evaluation of consistency of the updated model have been provided in Supplementary Text section S2.1 and Table S2. The reproducibility was verified for models of CBZ_IC50 and Control, with their respective FROG assessment reports provided in the supplementary. Further, as detailed in Supplementary Text section S2.2, with only the leaf module, the prediction of updated tomato leaf GEM in determining the trend in biomass decrease in the presence of nitrogen limitation was in alignment to VYTOP prediction following a similar pattern as reported experimentally by Groot et al. (2002). This confirms the updated model robustness in predicting the trend in specific growth rate under stress conditions.

**Figure 2:**
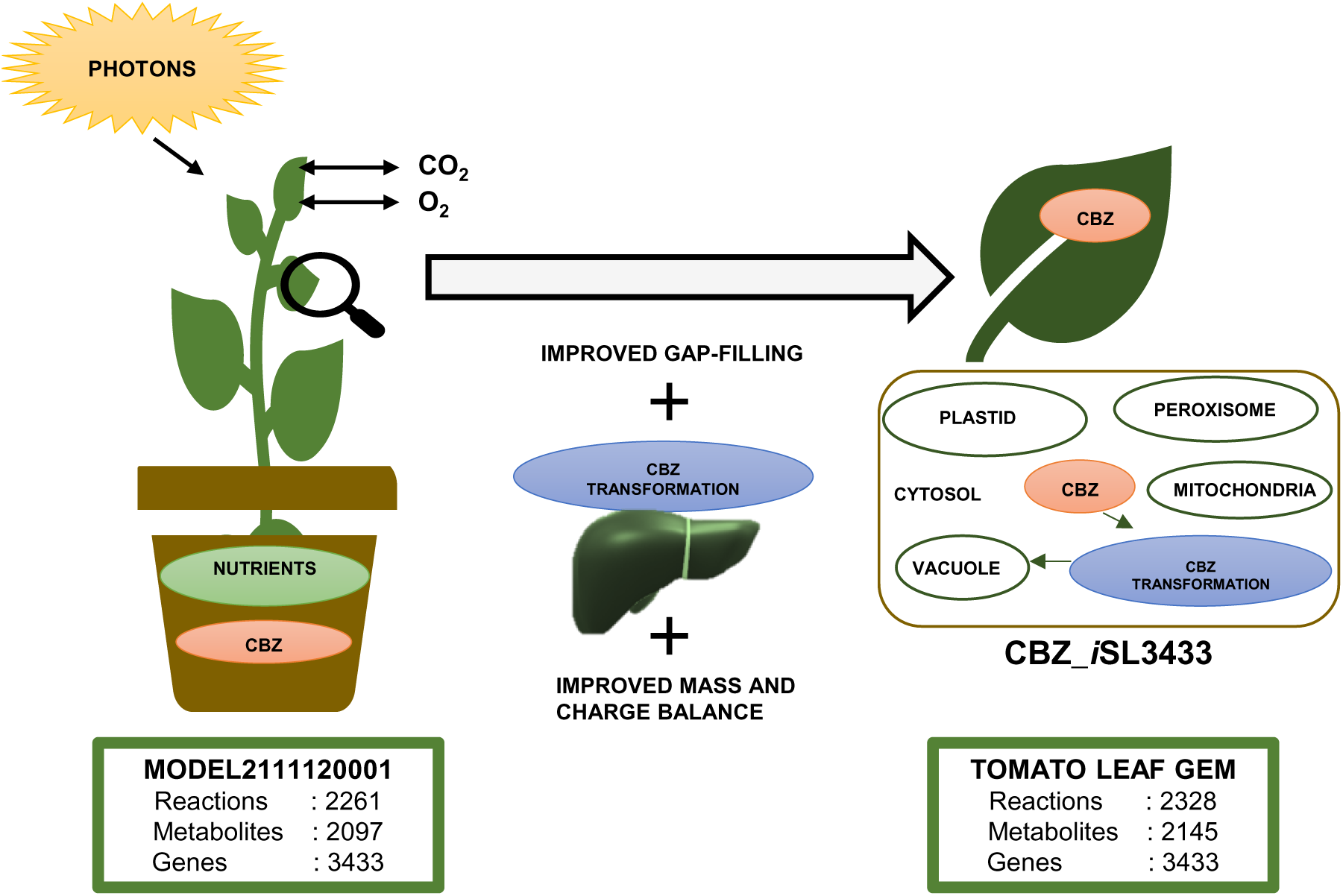
**Overview of steps in the development of updated tomato leaf GEM integrated with CBZ_PK module.**

### 3.2 Energy and Co-factor competition induced biomass reduction upon enforced CBZ uptake

In this study, CBZ uptake was enforced by fixing the upper and lower bounds of its exchange reaction to understand how plant metabolism is altered in response to its detoxification, as it is now its prerequisite for survival. The maximum possible specific growth rate was estimated for increasing CBZ uptake rates with constraints as per Table S1. As observed in Figure 3, the specific growth rate was found to be significantly decreased with increasing CBZ uptake rates. This observation was in accordance with literature wherein CBZ has been reported to hinder the growth of several plant species such as zucchini (Carter et al. 2015), *Typha latifolia* (Dordio et al. 2011), lettuce (Leitão et al. 2021b) plants. Furthermore, a concentration of 1000 ppb of CBZ-containing psychoactive pharmaceutical cocktail was found to be highly detrimental to tomato plants, reportedly resulting in plant collapse (Gorovits et al. 2020). Further, the simulation was repeated with biomass reaction upper bound value of 1 h^-1^ to evaluate the effect of constrained biomass value in model prediction of the impact of CBZ. As observed from Figure 3, a decrease in specific growth rate in the presence of increasing CBZ uptake rates was estimated to be independent of the initial constrained upper bound value of biomass except with a difference in the rate of decrease between the two conditions.

**Figure 3:**
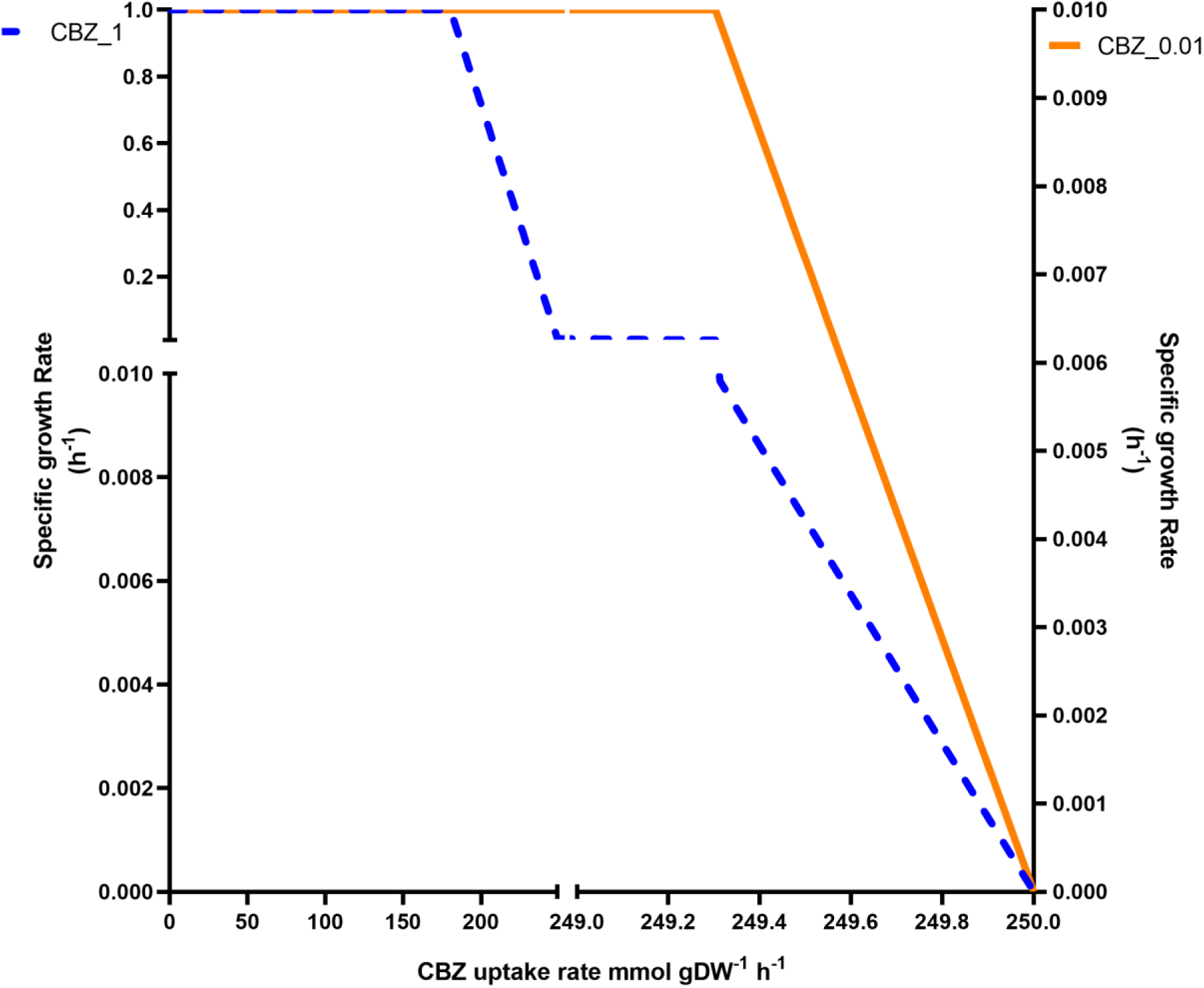
**Effect of increasing enforced uptake rates of CBZ on specific growth rate. A) Straight line representing ‘CBZ_0.01’ with maximum biomass constrained value of 0.01 h^-1^. B) Dotted line representing ‘CBZ_1’ with maximum biomass constrained value of 1 h^-1^.**

Plants are sessile and have evolved a wide range of tolerance mechanisms to endure the forced fluctuations in their environment. Unlike microbial systems, plants majorly employ detoxification mechanisms to avoid the potential build-up of toxicity due to xenobiotics like CBZ, rather than relying on catabolism for their elimination. The interconnection between the two networks of updated base tomato leaf GEM vs CBZ_PK module is represented in Figure 4. CBZ transformation is facilitated via utilization of metabolites like reduced form of Nicotinamide adenine dinucleotide phosphate (NADPH), Oxygen (O_2_), proton (H+), water (H_2_O), Uridine diphosphate glucose (UDP-Glucose), Pumped-proton (energetic proton representing ATP synthesis driven metabolic transfer across compartments) and Glutathione. Thus, tomato metabolism requires an additional expenditure of these metabolites in the presence of CBZ, thereby posing a metabolic burden to the cell, disrupting its homeostasis and inducing stress. This was also observed in drug-induced toxicity in the animal liver system, wherein the induced demand of drug detoxification has been observed to perturb cellular homeostasis (Cordes et al. 2018).

**Figure 4:**
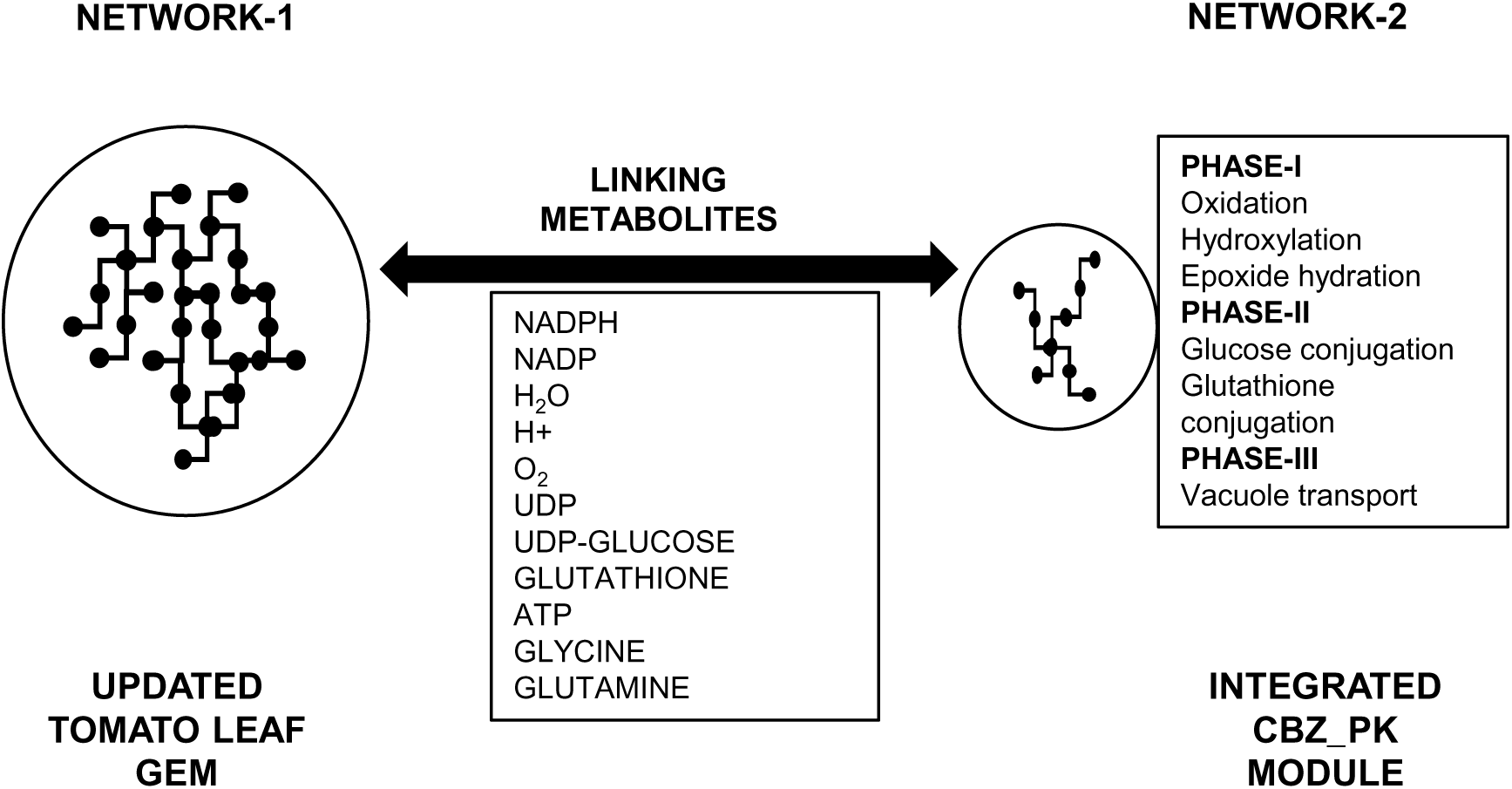
**Representation of linking metabolites between two networks of updated tomato leaf GEM and the integrated CBZ_PK module.**

### 3.3 CBZ uptake is predicted to elicit abiotic stress response in plants

Plants have been known to have evolved perfect metabolic plasticity to sustain challenging conditions with robust solutions for a sessile lifestyle (Knudsen et al. 2018). Pharmaceuticals fall into the category of toxic chemicals for plants as they are not able to be used for nutrition or source of energy (Coleman et al. 1997). Owing to their sessile nature, plants have evolved mechanisms to take up these chemicals and detoxify them. Xenobiotics are known to be chemically transformed to a more hydrophilic state to restrict their interaction with biological molecules and are further dumped in the vacuole or cell wall. To orchestrate this dynamic response in nature, plants have evolved abundant linear and non-linear feedforward/feedback regulations at the enzyme level to maintain metabolic steady state (Knudsen et al. 2018; Shameer et al. 2019). This forms the core basis of the mathematical framework of constraint-based modelling, which employs linear optimization by maximizing/minimizing an objective function for the determination of steady-state flux distribution of reactions in a metabolic network (Varma and Palsson, 1994; Kauffman et al. 2003; Raman and Chandra 2009). In the GEM, the presence of CBZ was captured by enforced higher CBZ uptake rates. A biologically relevant standard measure of CBZ uptake rate inducing a 50 % reduction in the specific growth rate of 0.005 h^-1^ from experimentally reported value under control conditions of 0.01 h^-1^ was chosen as the benchmark reference point to mimic the presence of CBZ stress for further simulations.

Assessing biomass is a widely used method for measuring stress response, as reduced growth represents the overall cumulative responses within the plant. To investigate the mechanistic insights on the predicted decrease in specific growth rate with increasing CBZ uptake rates, FVA and SI based framework was employed for identifying significantly altered metabolism. Approximately 31.3 % (728/2328) reactions had their SI in the range of 0 to 0.1 representing enhanced metabolic reprogramming in the presence of CBZ (Figure 5). Among these, 154 metabolic reactions were identified as significantly altered reactions with zero SI, indicating non-overlapping flux spaces between Control vs CBZ_IC50 conditions.

**Figure 5:**
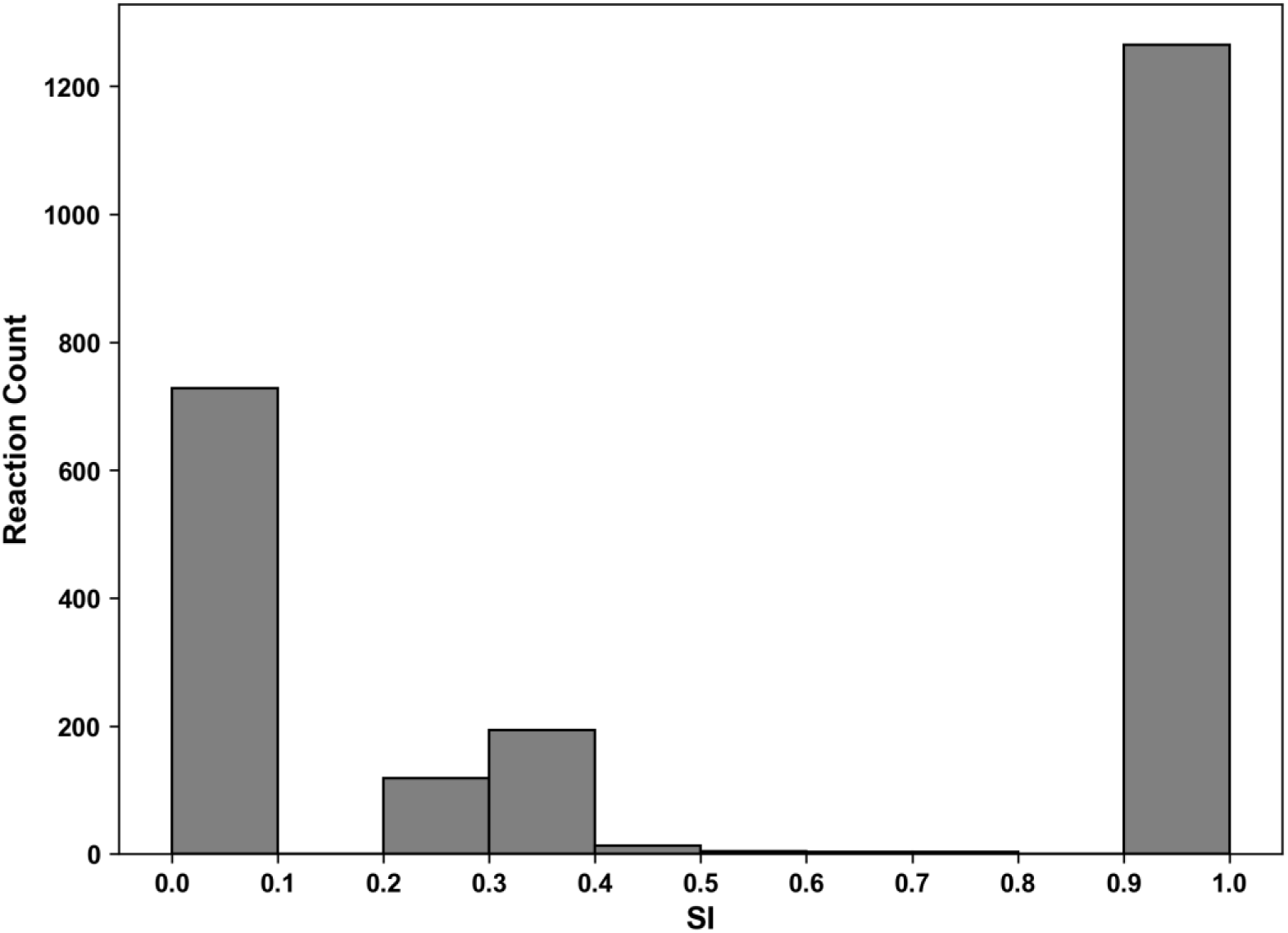
SI distribution of flux space of reactions in CBZ_*i*SL3433 compared between Control and CBZ_IC50 conditions with maximizing biomass production as objective function.

Further, the predicted list of altered reactions was verified to ensure that the predictions were independent of objective function choice and chosen CBZ uptake flux rate (Supplementary Text section S3).

### 3.4 Verification of CBZ Pharmacodynamics: *In silico* predictions vs literature data

The pathway classes of reactions representing significant alterations in the presence of CBZ is represented in Figure 6. The direct effect on biomass precursors was evident from the alterations in amino acid biosynthesis, chlorophyll biosynthesis, cell structure biosynthesis, purine nucleotide biosynthesis, fatty acid and lipid biosynthesis, ascorbate biosynthesis, carbohydrate biosynthesis and degradation, amino acid degradation and purine and pyrimidine nucleotide degradation. Also, methylerythritol phosphate pathway (MEP) and mevalonate pathway (MVA) involved in terpenoid and polyprenyl biosynthesis were also found to be significantly altered. Though directly are not a part of biomass components, the alterations predicted in the core pathways of pentose phosphate pathway, folate biosynthesis and transformation, photorespiration and Calvin cycle highlight the in-direct control of these pathways in biomass synthesis. Further, to investigate the perceived impact of CBZ in the network, the flux range of specific reactions in the major core pathway was compared between Control and CBZ_IC50 conditions. The flux ranges for reactions of major pathway classes are known to be altered under pharmaceutical stress shown in Figure S6, demonstrating CBZ-induced effect across all the pathways. 512 reactions were categorized into 20 pathways, with 37 reactions shared between multiple pathways.

**Figure 6:**
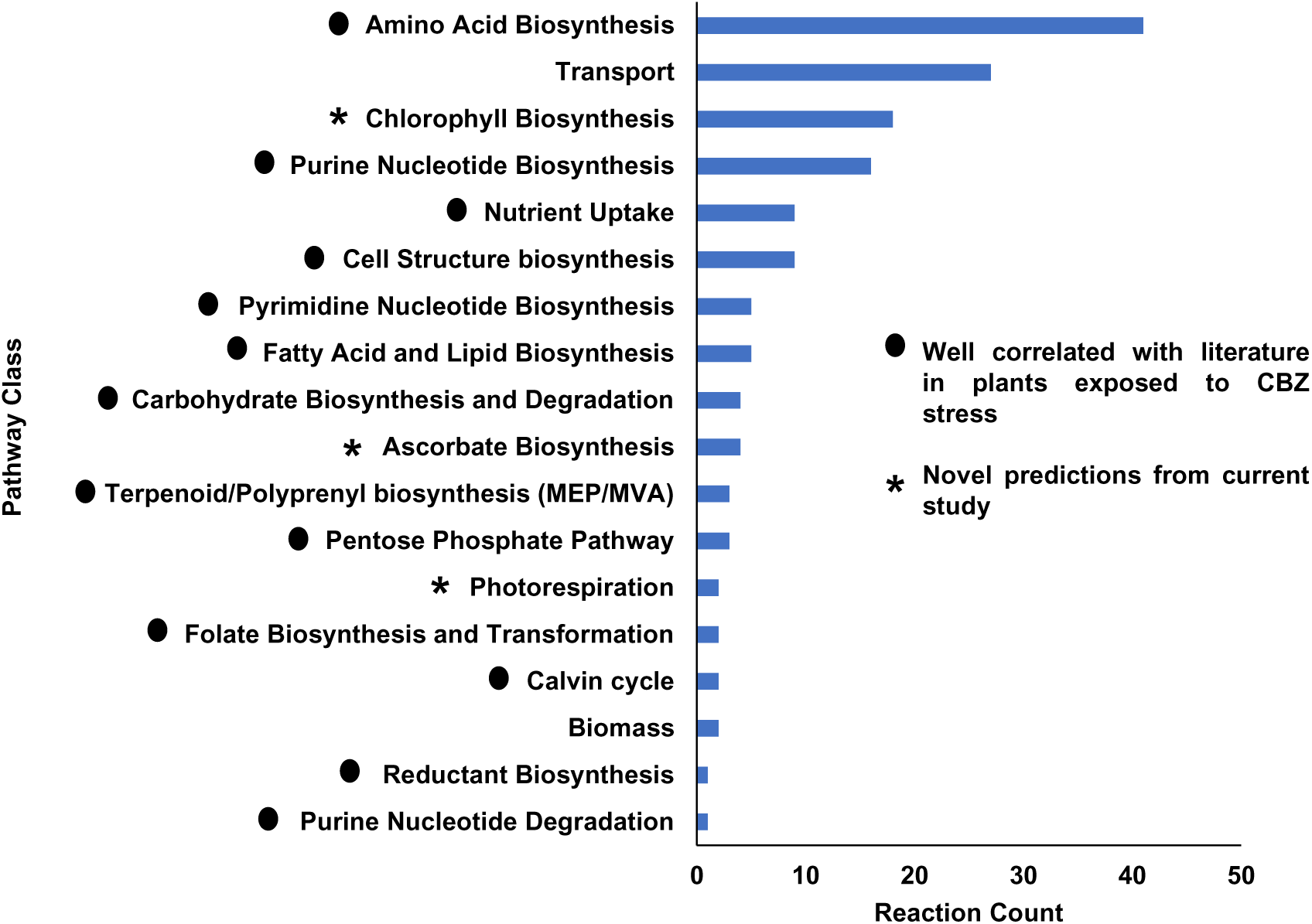
**Pathway classes of reactions (154) significantly altered in the presence of CBZ. The prediction of pathway is categorized as either consistent with existing literature or as novel predictions.**

Interestingly, the majority of the list of pathway classes altered with CBZ, represented in Figure 6 and Figure S6, were well correlated with literature based on CBZ exposure in plants (Gorovits et al. 2020; Wahman et al. 2020; Leitão et al. 2021a; Lu et al. 2024). Based on untargeted metabolomics in *Phragmites australis* plants, alterations in carbon fixation, pentose phosphate pathway, fatty acid biosynthesis, folate biosynthesis, purine and pyrimidine metabolism and amino acid metabolism (arginine, proline, tyrosine, β-Alanine, and tryptophan) has been reported in the presence of CBZ (Wahman et al. 2020). Further, in lettuce plants exposed to CBZ, alterations in oxidative stress response, cellular respiration, cell wall metabolism and phenylpropanoid metabolism have been observed (Leitão et al. 2021a).

Specifically in tomato plants, based on an integrated transcriptomics and metabolomics study, exposure to CBZ has been reported to increase ROS levels with induction of spatial specific alterations in leaf edges (Lu et al. 2024). This has been reported to alter antioxidant enzyme expression, photosynthesis, glutathione metabolism, starch and sucrose metabolism. Based on model predictions, alterations in glutathione metabolism, starch and sucrose metabolism were observed with SI for reactions in the pathway classes falling in the range of 0 to 1, demonstrating good correlation with experimental observations (Table S4). Also, altered synthesis of carotenoids, alkaloids, flavonoids, terpenes, and nucleotides have been experimentally observed (Lu et al. 2024). This observation was captured based on predictions, despite the model not representing the complete pathway of secondary metabolism. The model indicated alterations in key precursors of secondary metabolites. Chorismate biosynthesis was predicted to be altered, which is the key precursor for flavonoids. Additionally, aliphatic amino acid metabolism was predicted to be altered, which are the key precursor for alkaloids. Also, MVA/MEP pathway-mediated terpene biosynthesis was predicted to be altered, which is the key precursor for carotenoids. Further, Lu et al. (2024), verified the trend in the alteration of photosynthesis and antioxidant expression with quantitative polymerase chain reaction. The comparison between CBZ_*i*SL3433 predictions and experimental observations of transcript levels of these two pathways is represented in Table S3. CBZ_*i*SL3433 was able to capture the reduced photosynthesis but was not able to capture the increased antioxidant levels.

A prediction is considered robust when the correct pathway, along with the trend of increase/decrease is captured. Plant stress responses are species-specific and influenced by stressor dosage, timing, and environmental factors. This information can be mostly captured usually with transcriptomics or proteomics analysis for a particular instance. Thus, omics-integrated GEM have been successfully used for investigating stress response in plants (Lakshmanan et al. 2016; Wanichthanarak et al. 2020; Chowdhury et al. 2023). As per a proteomics study in lettuce, based on molecular functions of differentially expressed proteins in leaves, CBZ has been reported to induce 38.5 % of binding, 42.3 % of catalytic, 7.7 % of molecular function regulator activity and 11.5 % of structural molecular activity (Leitão et al. 2021a). For nitrogen stress prediction in the maize root model, Chowdhury et al. (2022) reported that, transcriptomics integration to GEM improved the robustness of model prediction with true prediction of metabolite fold change improved from 18 % to 70 % and false prediction rate reduced from 82 % to 30 %. Thus, this explains the discrepancy of model prediction for antioxidant expression, which can be improved with the integration of regulatory constraints in the model.

Further alterations in chlorophyll biosynthesis, ascorbate biosynthesis, photorespiration and nutrient assimilation of phototrophic input were predicted with CBZ presence. Chronic exposure of pharmaceuticals is known to have the potential to induce alterations in ammonium and nitrite assimilation (Leitão et al. 2021a). In *Cucurbita pepo* plants, reduction in chlorophyll pigments and increased nutrient composition, including phosphorous, potassium, magnesium, silicon, zinc, manganese and total nitrogen of leaves has been reported in the presence of CBZ (Carter et al. 2015; Knight et al. 2018). In lettuce plants exposed to pharmaceuticals, alterations in sulphur assimilation have been observed (Leitão et al. 2021a). Ascorbate biosynthesis and photorespiration being key-players of abiotic stress response and tolerance, are known to aid in redox homeostasis (Voss et al. 2013; Akram et al. 2017). Alterations predicted by the model for these pathways are novel predictions under CBZ stress response in plants which needs further validation. Thus, this highlights the potential of network-level approach to understand the metabolic perturbations. Based on transcriptomics analysis, glycerophospholipid, alpha-linolenic acid and linoleic metabolism were reported to be altered in tomato plants exposed to CBZ (Lu et al. 2024). Though model predicted altered lipid metabolism, these specific pathways are gaps in the model highlighting the future scope for improvement in the model.

### 3.5 *In silico* proof for reduced toxicity of CBZ in presence of biostimulants

Under challenging conditions of (a)biotic stress, plants have developed on-demand synthesis of array of phytochemicals to protect cells from injury induced from stress (Knudsen et al. 2018). Exogenous application of these metabolites has been reported as an economical approach to alleviate abiotic stress response. The effect of these molecules has not yet been investigated for pharmaceutical stress. Furthermore, GEM has been successfully used for screening nutrient supplements to improve the bioremediation of herbicides in microbial systems (Ofaim et al. 2020; Dhakar et al. 2021). However, its potential use in plant systems for prediction of stress mitigation strategies is limited.

In this study, the effect of four biostimulants including proline, spermine, ethanol and glycerol, was investigated with respect to CBZ stress. With maximum biomass production as objective, based on FBA (*l*1-norm) analysis, the effect of increasing flux of biostimulant, on specific growth rate under CBZ_IC50 was evaluated. In the presence of CBZ, all of the biostimulants under investigation demonstrated increased specific growth rate as observed in Figure 7 and were predicted as non-toxic under control conditions (Figure S7). Notably, when the biostimulant flux rate was set beyond 0.07 mmol gDW^-1^ h^-1^ with any of the biostimulants, the effect of CBZ was found to be negligible. Hence, this flux rate was chosen to compare the effects among the biostimulants for ameliorating CBZ stress.

**Figure 7:**
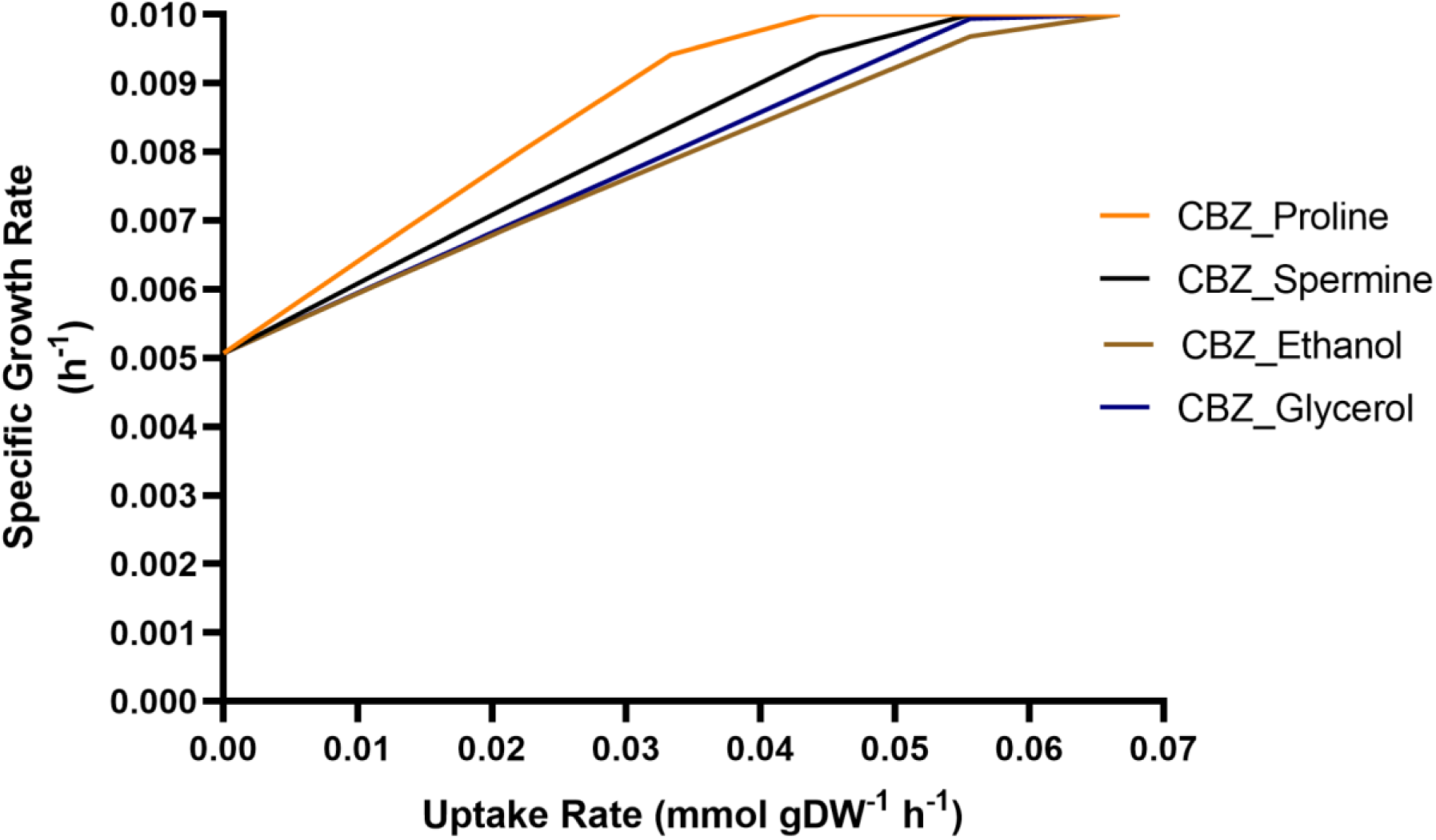
**Effect of exogenous addition of biostimulants (Proline, Spermine, Ethanol and Glycerol) on specific growth rate under CBZ_IC50 condition.**

This was in accordance with literature wherein biostimulants have been reported to improve plant growth under stress. Proline, an osmolyte and energy supplier, is known to maintain the NADPH:NADP+ ratio. In tomato plants exposed to salinity stress exogenous application of proline has known to improve growth (Kahlaoui et al. 2014). Spermine, a tetramine is known for prophylactic effects under stress conditions. In tomato plants, exogenous spermine has been known to improve plant growth under combined salinity and paraquat tolerance (Pascual et al. 2023). Ethanol has been reported to maintain cell energy status under stress conditions. In tomato plants exposed to heat stress, exogenous ethanol aided in alleviating heat damage by supporting reproductive development and vegetative growth (Todaka et al. 2024). Glycerol, sugar alcohol with pronounced osmolytic function has been known to improve plant growth under stress. For instance, improved growth was demonstrated under salt stress in *Ricinus communis* L. (M. Ali et al. 2008), maize (Kaya et al. 2013) and pistachio (Raoufi et al. 2020) with exogenous application of glycerol.

Further, as observed from Figure 8, the presence of biostimulants aided in significant enhancement of individual biomass components. Being providers of energy and redox carriers, biostimulants were able to thus aid in the additional energy and co-factor requirement for CBZ detoxification, thus improving biomass growth in the presence of CBZ. These predictions align with literature wherein exogenous proline has been reported to improve chlorophyll, total soluble carbohydrates, total free amino acids and ascorbic acid under varied abiotic stress (Hosseinifard et al. 2022). Similarly, spermine has been reported to upregulate accumulation of aconitic acid, gallic acid, sugars, amino acids under salt stress (Li et al. 2022). Further, under drought stress, ethanol addition has been known to induce osmolytes such as free amino acids and sugars (Bashir et al. 2022). Also, glycerol application has been known to improve chlorophyll under salt stress (Raoufi et al. 2020).

**Figure 8:**
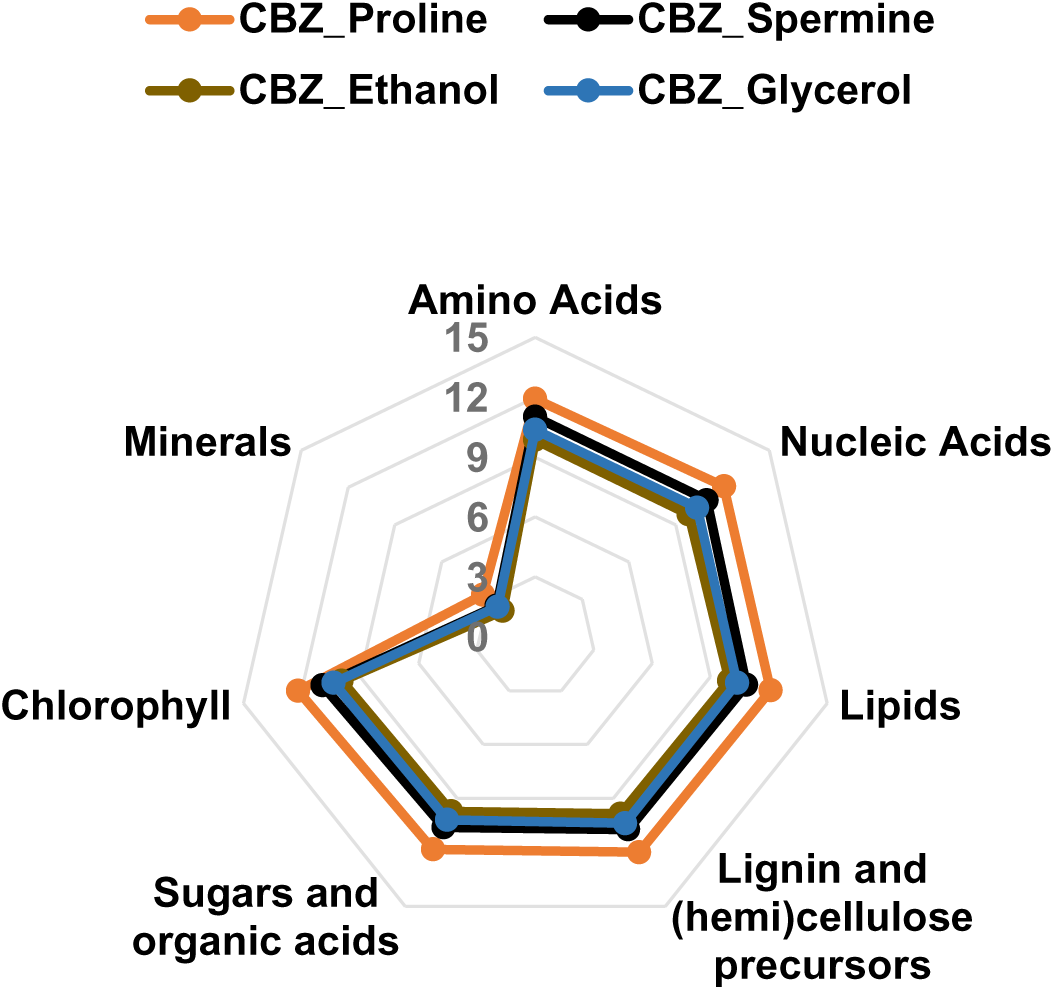
**Fold change ((CBZ_IC50_Biostimulant) / (CBZ_IC50)) representing production capacity of individual biomass components with exogenous addition of biostimulants (0.07 mmol gDW^-1^ h^-1^). Fold-change values were transformed into logarithm base 2.**

Interestingly, significant increase in phase-II and phase-III transformed metabolites of CBZ were observed in the presence of biostimulants (Figure 9). This could be due to increased availability of precursors of phase-II and phase-III mediated transformation in the presence of biostimulants. As observed from Figure 9, among biostimulants, proline induced higher fold changes, and this could be due to increased production capacity of UDP-Glucose and glutathione in the presence of proline (Figure S8). This was corroborated based on reports of enhanced xenobiotic detoxifying enzymes and improved glutathione with biostimulant exogenous addition under abiotic stress. Improved expression of genes of Cytochrome-P450 family, UDP-glycosyltransferases under drought stress (Bashir et al. 2022) and GST under salt stress (Nguyen et al. 2017) in *Arabidopsis* has been reported with ethanol addition. Also, improved GST activity in soybean under salt stress (Das et al. 2022) and drought stress (Rahman et al. 2022) have been reported with ethanol addition. Further, with spermine addition, improved GST lambda-1 protein was reported in oats under alkali stress (Bai et al. 2021). Further, spermine addition improved GST activity and glutathione content in *Vigna radiata* L. under cadmium stress (Nahar et al. 2016). Enhanced expression of genes of UDP-glycosyltransferases has been reported with glycerol addition under powdery Mildew disease in wheat (Li et al. 2020). Also, under stress, proline is known to be positively associated with glutathione and proline addition has been known to improve GST activity (Raza et al. 2023).

**Figure 9:**
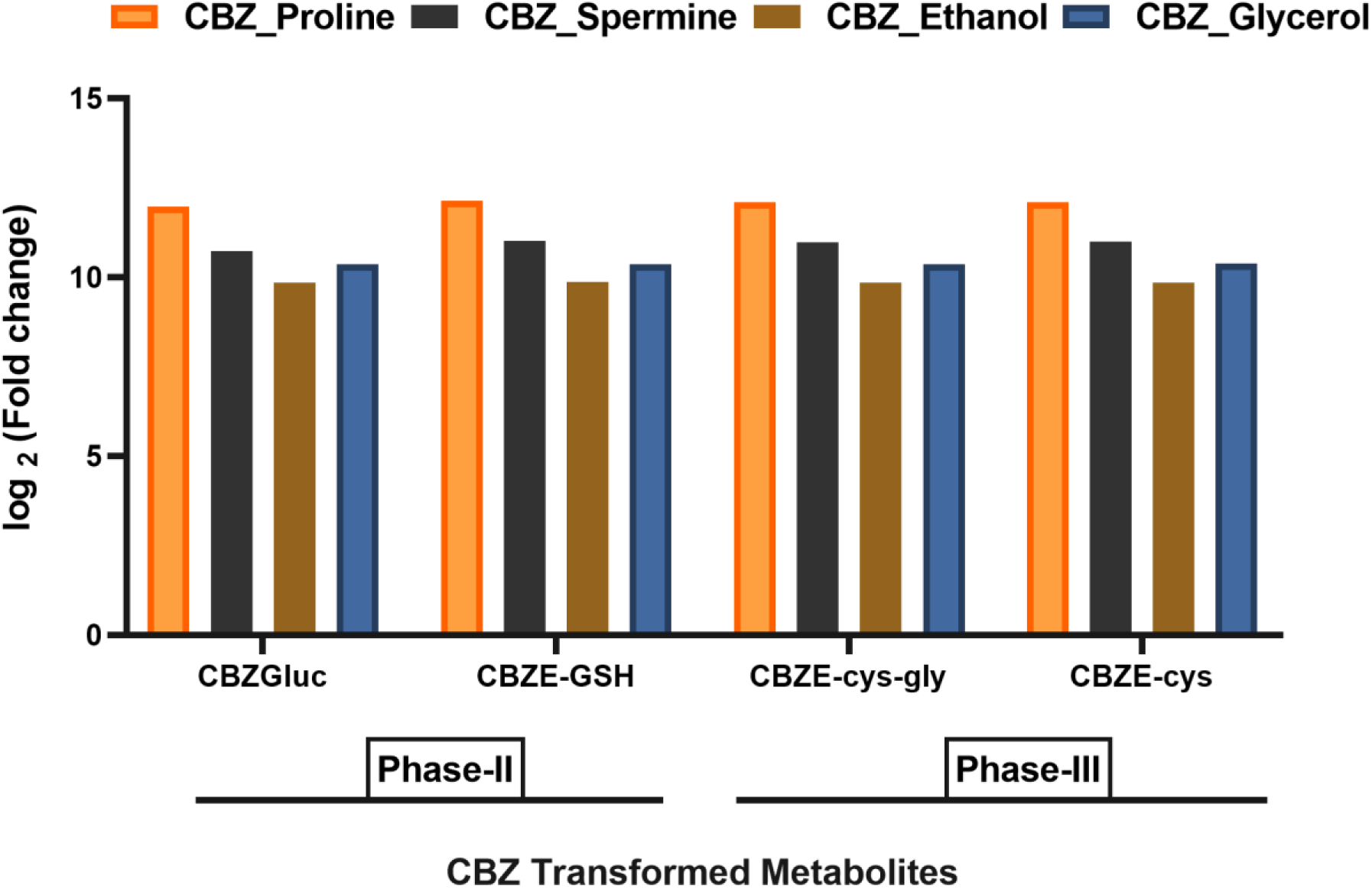
**Fold change ((CBZ_IC50_Biostimulant) / (CBZ_IC50)) of production capacity of CBZ transformed metabolites of Phase-II and Phase-III metabolism with exogenous addition of 0.07 mmol gDW^-1^ h^-1^ of biostimulants. Fold-change values were transformed into logarithm base 2.**

Interestingly, in animal liver system, biomolecules like sugar conjugation mediated phase-II transition have been known to reduce the toxicity of the xenobiotics making it more polar to be excreted from the system. Also, in context of plants, phase-II metabolites are reported as non-toxic or less phytotoxic molecules than its corresponding phase-I metabolites, with phase-II being an important protective phase in detoxification of xenobiotics (Coleman et al. 1997; Riemenschneider et al. 2016). Thus, in this study, the model predicted reduced toxicity of CBZ via increased detoxification in the presence of biostimulants is a novel prediction and warrants further experimental investigation. This prediction can be also be employed for facilitating easier analytical detection and quantification of phase-II and phase-III metabolites of pharmaceuticals like CBZ in the presence of biostimulants.

### 3.6 *In silico* proof for ameliorative role of biostimulants for CBZ stress in tomato

To comprehensively investigate the perceived impact of biostimulants under CBZ stress, FVA and SI based framework was employed for identifying significantly altered metabolism under CBZ_IC50 condition in presence and absence of biostimulants. Figures 10a, 10b, 10c and 10d, represent the effect of biostimulants on major pathways with proline, spermine, ethanol, and glycerol, respectively. 520 reactions were categorized into 21 pathways with 37 reactions shared between multiple pathways. It was observed that, biostimulants addition enhances significantly all the major pathways, thus improving biomass precursors production.

**Figure 10:**
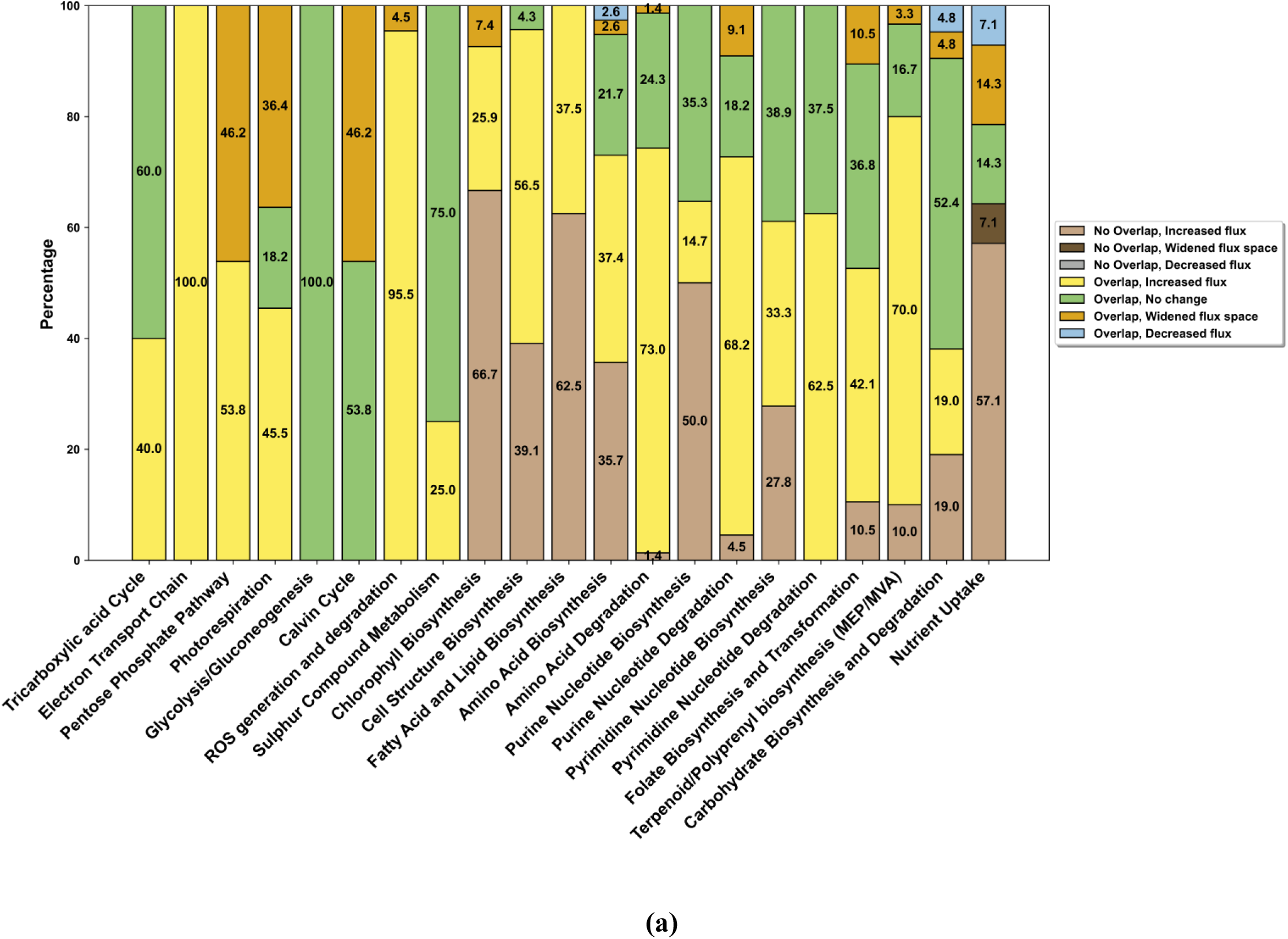
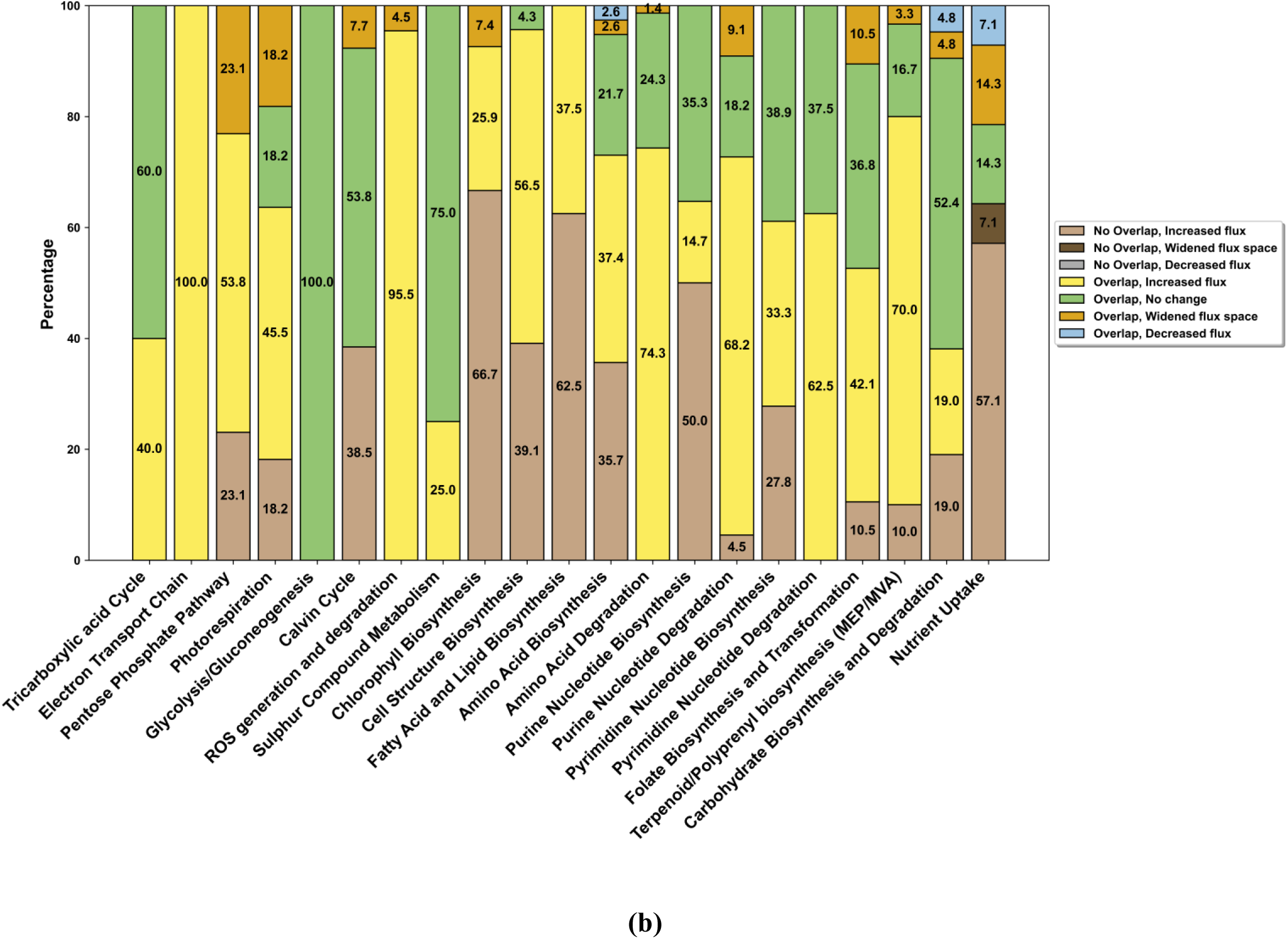
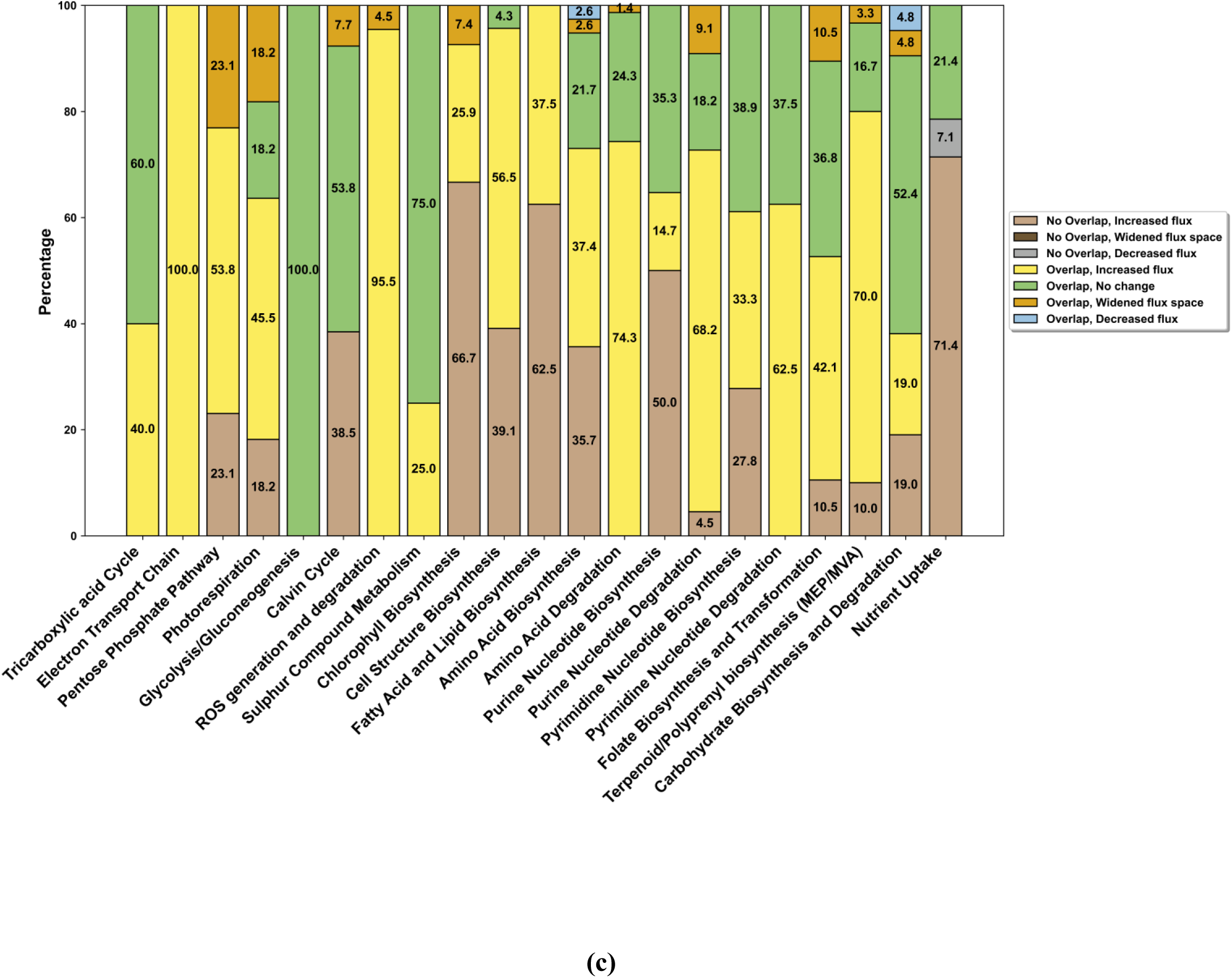
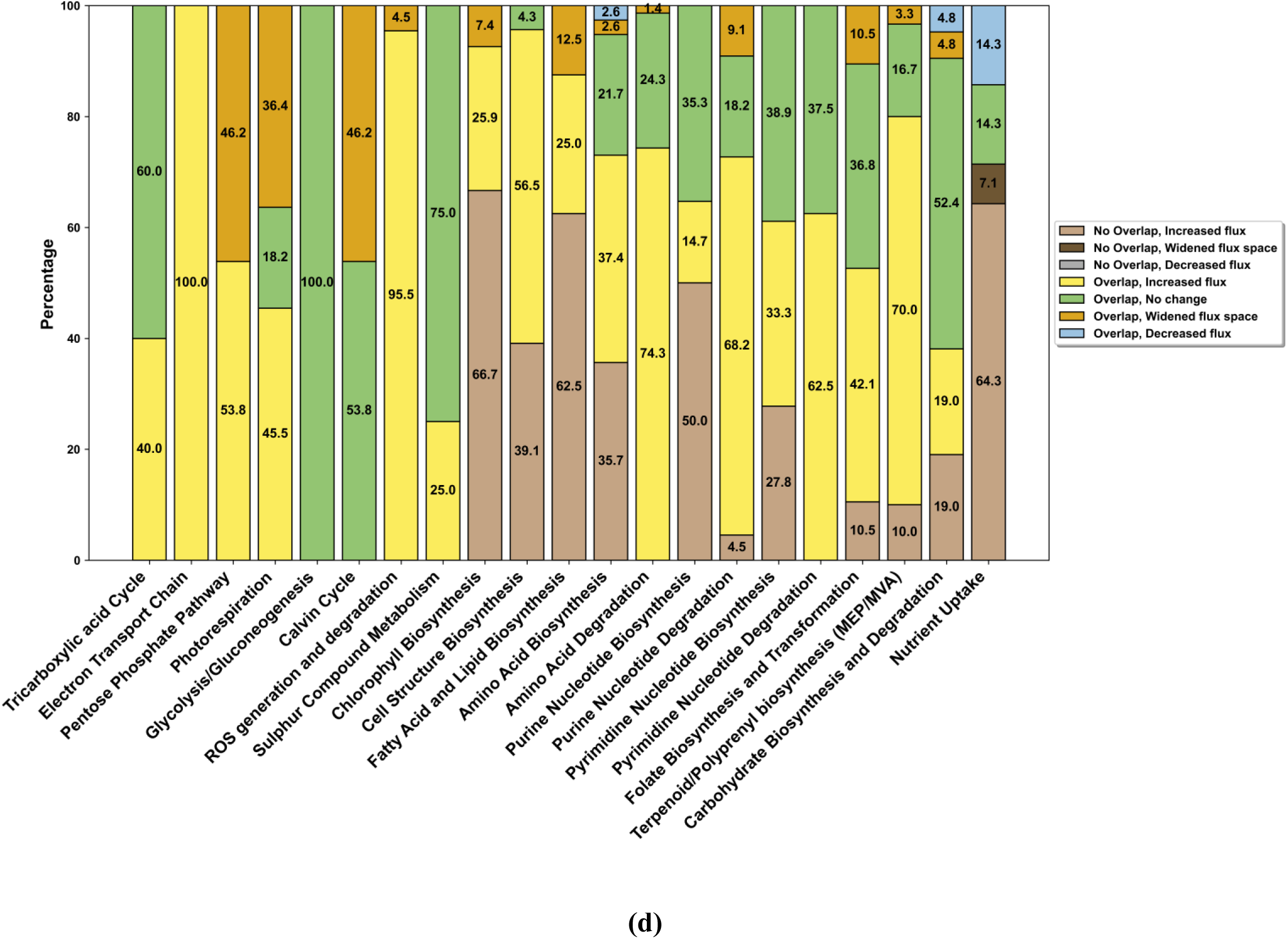
**A representation of altered metabolism in presence of biostimulants. Each bar-chart represents reprogramming of a specific metabolic pathway class in presence of biostimulants (a: Proline, b: Spermine, c: Ethanol; d: Glycerol) under CBZ stress. Numbers represent percentage of altered reactions in a pathway class. Each component of bar-chart represents one of the categories as mentioned in legend. Each category represents the classification of a reaction based on FVA analysis under CBZ_IC50 with different flux rates of biostimulants (0, 0.07 mmol gDW^-1^ h^-1^).**

Interestingly, ∼90-96.6 % of the significantly altered reactions in the presence of CBZ between the biostimulants were found to be in common, as represented in Figure 11a. This could be likely due to interplay among biostimulants, where their overlapping effects have been known to aid in fine-tuning adaptations to environmental cues. For instance, in *Agrostis stolonifera* plants under salt stress, exogenous spermine, improved glycerol (Li et al. 2022). Further, in switchgrass, exogenous proline was reported to improve polyamines metabolisms under salt stress (Guan et al. 2020). Also, with ethanol treatment in *Arabidopsis*, spermidine and glycerol was detected in response to drought stress (Bashir et al. 2022). Further, glycerol treatment was known to alter proline under salt stress in maize (Kaya et al. 2013).

**Figure 11:**
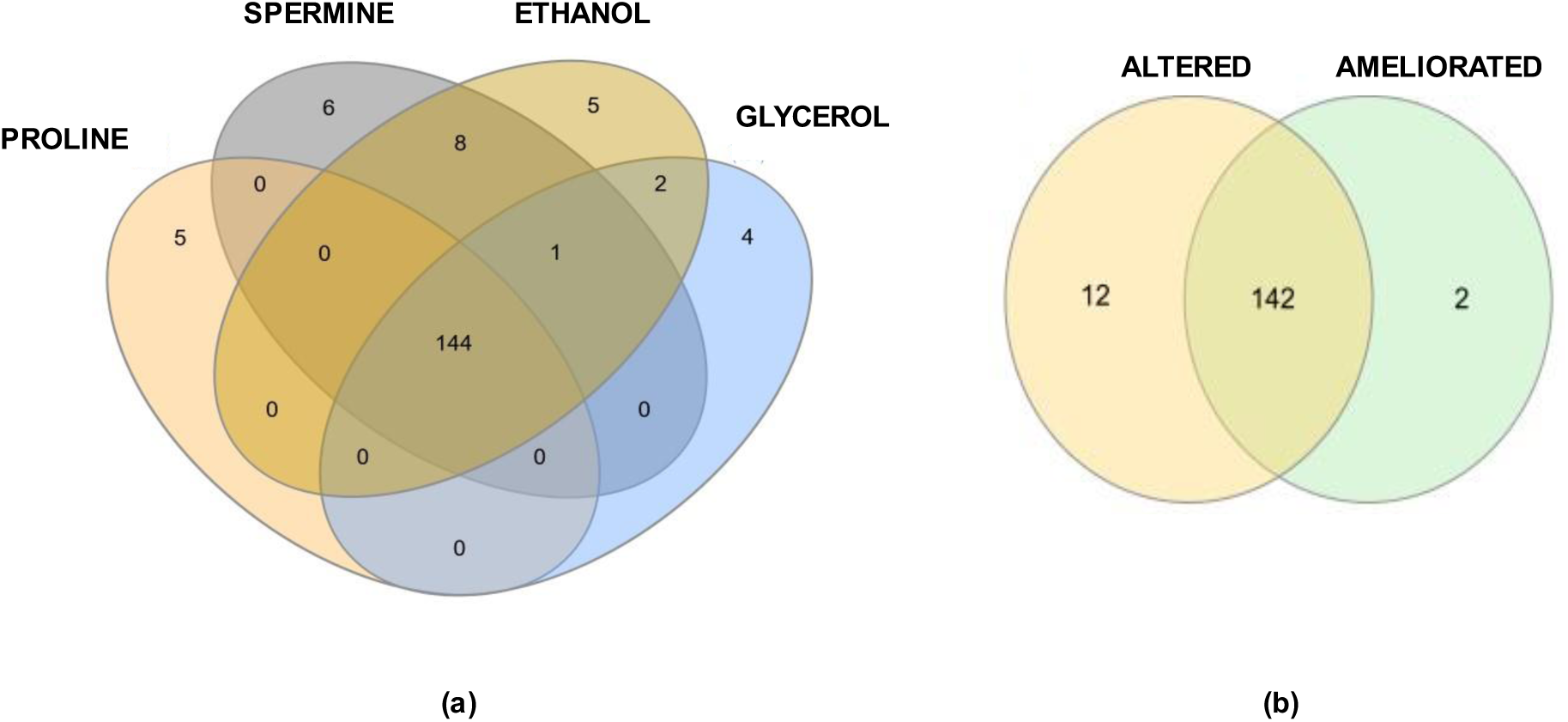
**Venn-diagram representation (Heberle et al. 2015). (a). Common significantly altered reactions between biostimulants. Zero SI reactions sets based on FVA results of CBZ_IC50 with individual biostimulants are compared. (b). Intersection of significantly altered reaction under CBZ and its amelioration with biostimulants. ‘ALTERED’ representing 154 zero SI reactions of FVA between Control and CBZ_IC50. ‘AMELIORATED’ representing 144 zero SI reactions commonly altered with biostimulants.**

Thus, in the current study, 144 reactions were found to be in common among the significantly altered reactions with biostimulants in the presence of CBZ. Notably, 98 % of common significantly altered reactions are a subset of 154 reactions known to be altered under CBZ stress, as mentioned in Section 3.4 (Figure 11b). Also, each of the primary breakdown step of the biostimulant was predicted among the uniquely altered reactions with respect to each biostimulant. Reactions including PROD2m (Proline dehydrogenase, EC Number:1.5.99.8), SMOX (Spermine oxidase, EC Number: 1.5.3.16), ALCD2x (alcohol dehydrogenase, EC Number: 1.1.1.1), GLYK (Glycerol kinase, EC Number: 2.7.1.30) representing the major breakdown of proline, spermine, ethanol and glycerol to pyrroline 5-carboxylate, spermidine & 3-aminopropionaldehyde, acetaldehyde, glyceraldehyhe-3-phosphate respectively was predicted as uniquely altered with respect to each biostimulant.

Further, in Figure 12, the pathway classes for the 144 reactions significantly altered are represented and among them the overview of metabolic alterations is depicted in Figure 13. Based on model predictions, biostimulants significantly improved the transformation of ribose-5-phosphate to 5-Phospho-α-D-ribose 1-diphosphate (PRPP) conversion subsequently improving histidine and purine metabolism. Improved orotate with improved PRPP led to improved Uridine monophosphate. Chorismate biosynthesis was enhanced, which improved tryptophan biosynthesis and phenylpropanoid mediated cell structure biosynthesis. Improved xylan biosynthesis via UDP-xylose was predicted. Also, mannose-6-phosphate mediated ascorbate biosynthesis was improved. Improved conversion of acetyl-CoA to fatty acids was predicted. Pyruvate mediated biosynthesis of isoleucine, valine and lysine was significantly enhanced. Nutrient assimilation and glutamate-mediated conversion of Chlorophyll-b was improved. Further, Deoxyuridine monophosphate mediated conversion to dihydrofolate and tetrahydrofolate was enhanced. Also, terpenoid synthesis via MEP pathway, with synthesis of geranyl phosphate, farnesyl phosphate, geranyl diphosphate production was also found to be improved.

**Figure 12:**
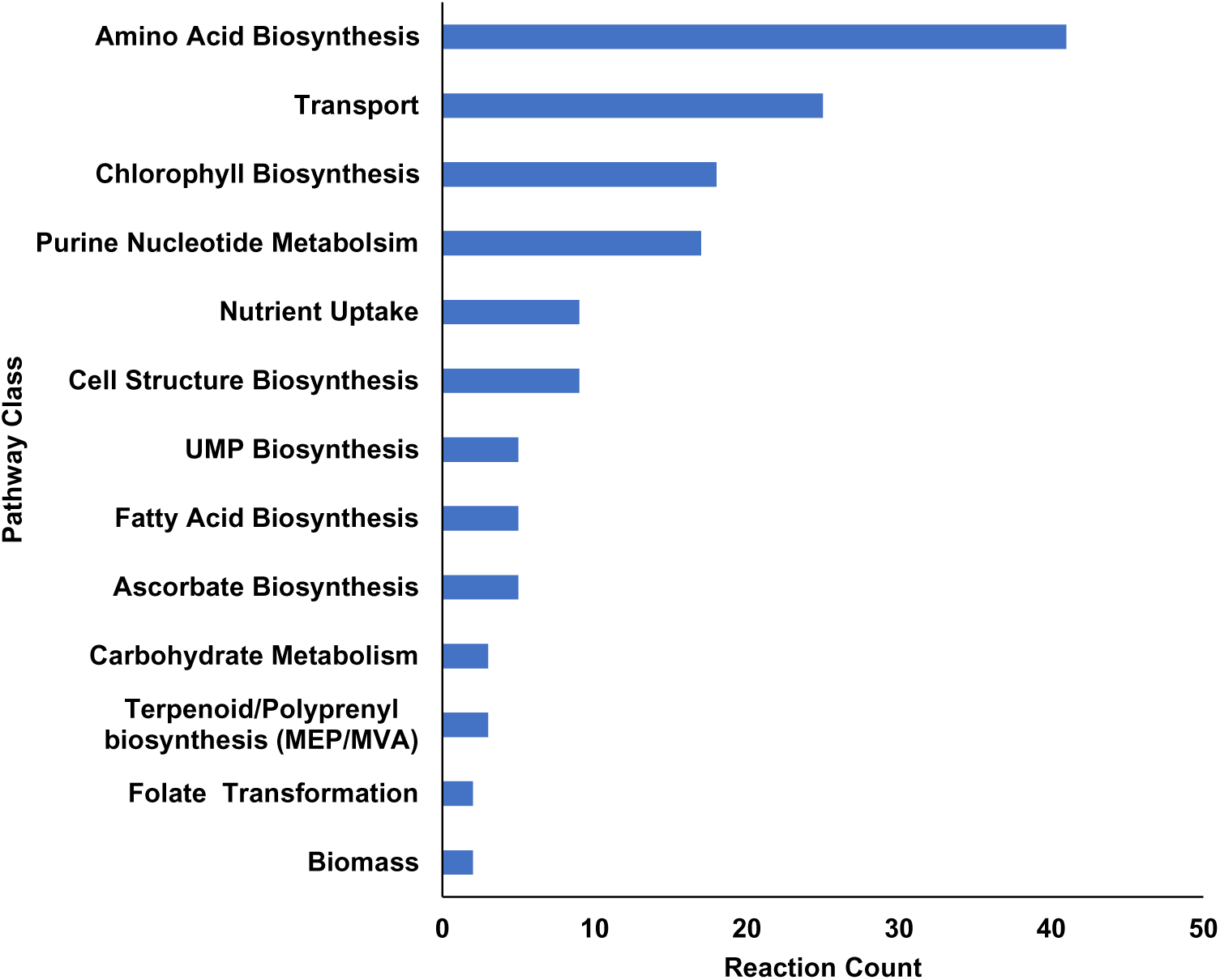
**Pathway classes for reactions (144) significantly altered in presence of biostimulants under CBZ stress.**

**Figure 13:**
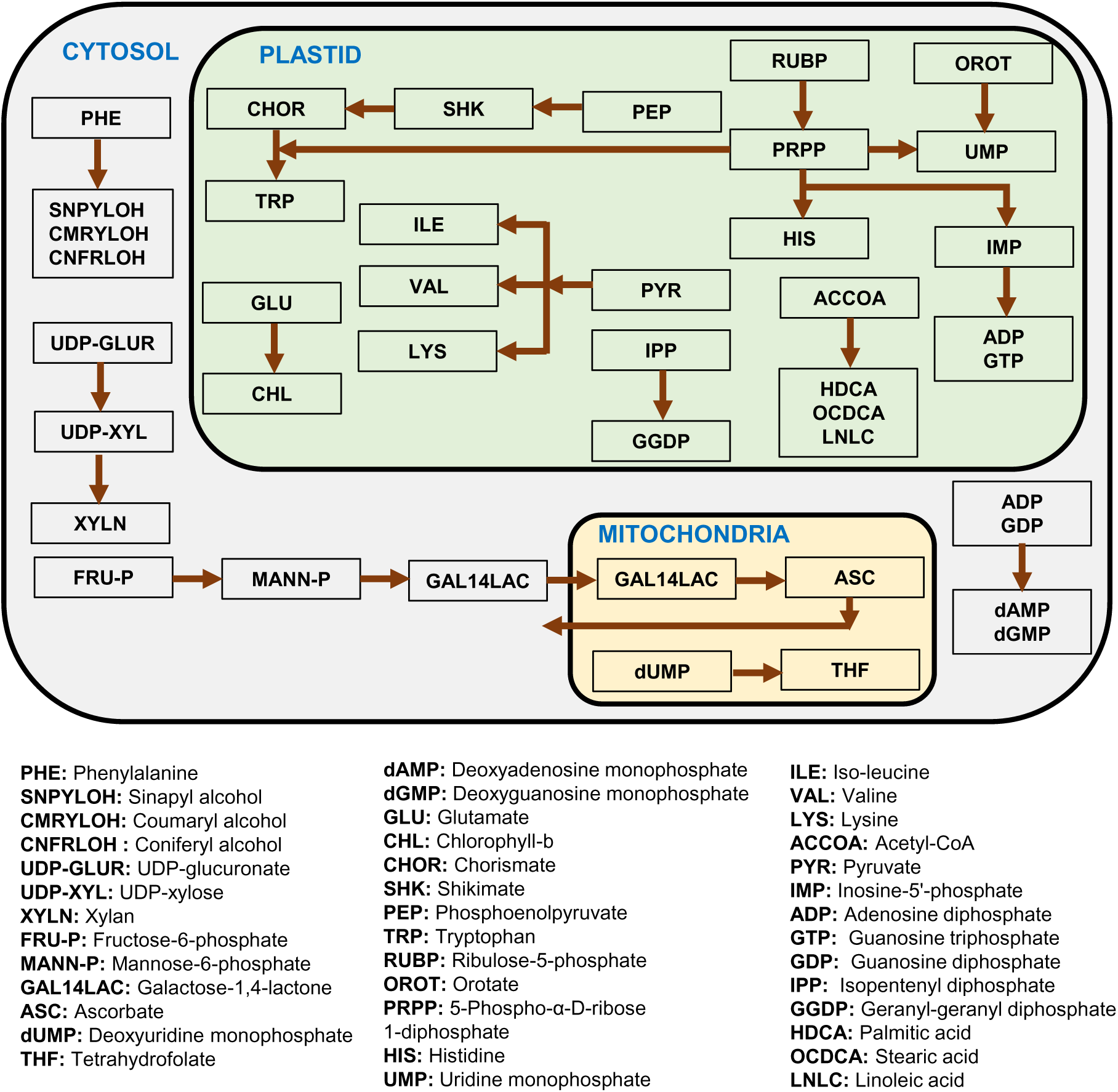
**An overview of predicted significantly altered metabolic reactions impacted with CBZ stress and ameliorated with biostimulants.**

This was consistent with trends observed with biostimulants in alleviating other stress. Under various stress conditions, proline has been reported to aid in improved nutrient uptake, photosynthesis and antioxidant enzyme activity (Hosseinifard et al. 2022). Spermine is known to improve photosynthesis, carbohydrate metabolism, antioxidant capacity, Krebs cycle, Gamma-aminobutyric acid (GABA) shunt, glycolysis, and nucleotide metabolism with exogenous application under abiotic stress (Bai et al. 2021; Hasan et al. 2021; Li et al. 2022).

Exogenous addition of ethanol has been known to improve nutrient uptake, photosynthesis, photorespiration, sucrose/starch metabolism, phenylpropanoids biosynthesis, glycolysis, fermentation, amino-acid biosynthesis, antioxidant defence response, gluconeogenesis under various abiotic stress (Bashir et al. 2022; Das et al. 2022; Todaka et al. 2024). Glycerol has been reported to improve nutrient uptake, chlorophyll, fatty acid metabolism under (a)biotic stress (M. Ali et al. 2008; Li et al. 2020; Raoufi et al. 2020).

Key metabolites have been known to be integral part of plant stress response. According to Chowdhury et al. (2023), reactions incorporated with transcriptomics constraints, are classified as bottleneck reactions of stress response if the expansion of their flux bounds improved biomass flux. In the current study, as detailed in Supplementary Text section S1.5, the same analysis was adapted for exchange reactions to screen for metabolites, which upon exogenous addition alleviates CBZ stress. Based on model prediction, 12 metabolites spanning major components of core pathways and 36 direct biomass precursors were identified as key metabolites for CBZ stress response. Some of these metabolites as compatible osmolytes have been known to accumulate as a part of stress adaptation in plants wherein in tomato plants increased proline, sucrose, glucose, fructose, leucine, isoleucine, valine, and GABA were observed under CBZ stress (Gorovits et al. 2020, 2021). In the current study, CBZ induced energy and co-factor demand reduced availability of these key metabolites. Further, biostimulants improved the availability of these key metabolites of core pathway (Figure S9) and subsequently the biomass precursors (Figure 8), thus aiding in improved biomass under CBZ stress. Thus, with the GEM serving as a screening tool, this approach can be further extended to explore plant-microbial interactions for alleviating CBZ stress and screen for more cost-effective biostimulants as alternatives that induce responses similar to those of the biostimulants used in this study by increasing the availability of these key metabolites. The list of key metabolites and significantly altered reactions are provided in the supplementary file ‘Supplementary_2.xlsx’. CBZ_*i*SL3433 predictions are based on stoichiometric model, thus model connectivity and composition of biomass equation play a crucial role in model predictions. The base model contains a few imbalanced reactions due to missing co-factor information which warrants future scope for improvement. This study thus highlights the pipeline providing first-guess type estimates for rational investigation of plant stress response to xenobiotics like pharmaceuticals which can be further validated and its accuracy improved by incorporating transcriptomic/proteomic data as regulatory constraints.

## 4. Conclusion

Innovative strategies have to be devised to achieve higher crop yields with diminishing land areas to meet the food security needs, especially in the face of climate change. By integrating the green-liver concept with metabolic modelling, this study provides a computational framework for understanding the metabolic impact of pharmaceutical pollutants on crops, offering a predictive tool for optimizing stress mitigation strategies.

An updated tomato leaf GEM was integrated with CBZ detoxification module. The developed model, CBZ_*i*SL3433 was able to capture the energy and co-factor mediated demand of CBZ stress leading to reduced biomass. Based on model predictions, 48 key metabolites and 154 significantly altered reactions were identified as a part of CBZ stress response. With a stoichiometric network, based on law of mass action and thermodynamics, model was able to capture the key altered metabolic reprogramming states in the presence of CBZ in core metabolism (Electron Transport Chain, Photorespiration, Photosynthesis, biomass precursors synthesis etc), synthesis of precursors of secondary metabolites (chorismate, terpenoid, phenylpropanoid) and nutrient assimilation reported in CBZ exposure in plants. Also, the model predicted the impact of the mitigation strategy with foliar addition of biostimulants (proline, spermine, ethanol, and glycerol) to combat CBZ stress. Biostimulants aided with improving biomass under CBZ stress and additionally improved phase-II and phase-III metabolism of CBZ.

The proposed framework serves as a screening tool prior to experimentation, thus aiding in reducing contaminant exposure by minimizing hit-and-trial experimental investigation. Further, the distribution of pharmaceutical mixtures in various parts of a plant can be influenced by their physiochemical properties, and the current model can be extended to build a robust whole-plant-based model for understanding the pharmaceutical mixture response. Physiological based pharmacokinetic modelling for pharmacokinetics, combined with integrated omics and diel cycle-based GEM will be a valuable tool in the way forward for plant pharmacovigilance.

## Author Contributions

Samyuktha Srinivasan: Methodology, Investigation, Data curation, Visualization, Writing – Original Draft.

Karthik Raman: Supervision, Conceptualization, Formal analysis, Validation, Writing – Review and editing.

Smita Srivastava: Supervision, Resources, Conceptualization, Formal analysis, Validation, Writing – Review and editing.

All authors read and approved the final manuscript.

## Supporting information

CBZ_iSL3433.xml

CBZ_iSL3433_CBZ_IC50.omex

CBZ_iSL3433_Control.omex

## Acknowledgement

Prof. Smita Srivastava would like to thank Hyclone Life Sciences Solutions India Private Limited (Project Number: CR22230114BTHLSS008458) and L&T Technology Services Limited (Project Number: CR21221810BTLNTE008458) under Corporate Social Responsibility (CSR) for funding the research work in Plant Cell technology Lab at Department of Biotechnology, Bhupat and Jyoti Mehta School of Biosciences, IIT Madras. Samyuktha Srinivasan would like to thank Dr. Maziya Ibrahim and Lavanya Raajaraam of Computational Systems Biology Lab, IIT Madras, for assistance regarding the use of COBRA Toolbox and relevant discussions to improve the associated analysis.

## Data Availability

All data supporting the finding of this study are available in the paper and its supplementary section.

## Supplementary Data

**Supplementary Text:** This supplementary document includes detailed methods, additional analysis, and supplementary figures that support the findings presented in the study.

**Supplementary_1.xlsx:** The updates to the base tomato model are summarised. The pathway categories for reactions investigated in this study are also provided in this file.

**Supplementary_2.xlsx:** The details of CBZ pharmacokinetic module are included in this file. The list of 48 key metabolites and 154 significantly altered reactions of CBZ stress is also included in this file.

**CBZ_*i*SL3433.xml:** The updated model file in SBML format is provided.

**CBZ_iSL3433_CBZ_IC50.omex:** FROG Report for the updated model under CBZ_IC50 condition is provided.

**CBZ_iSL3433_Control.omex:** FROG Report for the updated model under Control condition is provided.

## References

1. Akram, N.A., Shafiq, F., Ashraf, M., 2017. Ascorbic Acid-A Potential Oxidant Scavenger and Its Role in Plant Development and Abiotic Stress Tolerance. Frontiers in Plant Science, 8:613. 10.3389/fpls.2017.00613

2. Bai, J., Jin, K., Qin, W., Wang, Y., Yin, Q., 2021. Proteomic Responses to Alkali Stress in Oats and the Alleviatory Effects of Exogenous Spermine Application. Frontiers in Plant Science, 12: 627129. 10.3389/fpls.2021.627129

3. Bashir, K., Todaka, D., Rasheed, S., Matsui, A., Ahmad, Z., Sako, K., Utsumi, Y., Vu, A.T., Tanaka, M., Takahashi, S., Ishida, J., Tsuboi, Y., Watanabe, S., Kanno, Y., Ando, E., Shin, K.-C., Seito, M., Motegi, H., Sato, M., Li, R., Kikuchi, S., Fujita, M., Kusano, M., Kobayashi, M., Habu, Y., Nagano, A.J., Kawaura, K., Kikuchi, J., Saito, K., Hirai, M.Y., Seo, M., Shinozaki, K., Kinoshita, T., Seki, M., 2022. Ethanol-Mediated Novel Survival Strategy against Drought Stress in Plants. Plant and Cell Physiology 63, 1181– 1192. 10.1093/pcp/pcac114

4. Carter, L.J., Chefetz, B., Abdeen, Z., A. Boxall, A.B., 2019. Emerging investigator series: towards a framework for establishing the impacts of pharmaceuticals in wastewater irrigation systems on agro-ecosystems and human health. Environmental Science: Processes & Impacts 21, 605–622. 10.1039/C9EM00020H

5. Carter, L.J., Williams, M., Böttcher, C., Kookana, R.S., 2015. Uptake of Pharmaceuticals Influences Plant Development and Affects Nutrient and Hormone Homeostases. Environmental Science & Technology 49, 12509–12518. 10.1021/acs.est.5b03468

6. Chowdhury, N.B., Schroeder, W.L., Sarkar, D., Amiour, N., Quilleré, I., Hirel, B., Maranas, C.D., Saha, R., 2022. Dissecting the metabolic reprogramming of maize root under nitrogen-deficient stress conditions. Journal of Experimental Botany 73, 275–291. 10.1093/jxb/erab435

7. Chowdhury, N.B., Simons-Senftle, M., Decouard, B., Quillere, I., Rigault, M., Sajeevan, K.A., Acharya, B., Chowdhury, R., Hirel, B., Dellagi, A., Maranas, C., Saha, R., 2023. A multi-organ maize metabolic model connects temperature stress with energy production and reducing power generation. iScience 26, 108400. 10.1016/j.isci.2023.108400

8. Christou, A., Michael, C., Fatta-Kassinos, D., Fotopoulos, V., 2018. Can the pharmaceutically active compounds released in agroecosystems be considered as emerging plant stressors? Environment International 114, 360–364. 10.1016/j.envint.2018.03.003

9. Coleman, J., Blake-Kalff, M., Davies, E., 1997. Detoxification of xenobiotics by plants: chemical modification and vacuolar compartmentation. Trends in Plant Science 2, 144–151. 10.1016/S1360-1385(97)01019-4

10. Cordes, H., Thiel, C., Baier, V., Blank, L.M., Kuepfer, L., 2018. Integration of genome-scale metabolic networks into whole-body PBPK models shows phenotype-specific cases of drug-induced metabolic perturbation. NPJ Systems Biology and Applications 4, 10. 10.1038/s41540-018-0048-1

11. Das, A.K., Anik, T.R., Rahman, M.M., Keya, S.S., Islam, M.R., Rahman, M.A., Sultana, S., Ghosh, P.K., Khan, S., Ahamed, T., Ghosh, T.K., Tran, L.S.-P., Mostofa, M.G., 2022. Ethanol Treatment Enhances Physiological and Biochemical Responses to Mitigate Saline Toxicity in Soybean. Plants 11, 272. 10.3390/plants11030272

12. Dhakar, K., Zarecki, R., van Bommel, D., Knossow, N., Medina, S., Öztürk, B., Aly, R., Eizenberg, H., Ronen, Z., Freilich, S., 2021. Strategies for Enhancing *in vitro* Degradation of Linuron by *Variovorax* sp. Strain SRS 16 Under the Guidance of Metabolic Modeling. Frontiers in Bioengineering and Biotechnology, 9: 602464. 10.3389/fbioe.2021.602464

13. Dordio, A.V., Belo, M., Martins Teixeira, D., Palace Carvalho, A.J., Dias, C.M.B., Picó, Y., Pinto, A.P., 2011. Evaluation of carbamazepine uptake and metabolization by *Typha* spp., a plant with potential use in phytotreatment. Bioresource Technology 102, 7827–7834. 10.1016/j.biortech.2011.06.050

14. García-García, A.L., García-Machado, F.J., Borges, A.A., Morales-Sierra, S., Boto, A., Jiménez-Arias, D., 2020. Pure Organic Active Compounds Against Abiotic Stress: A Biostimulant Overview. Frontiers in Plant Science, 11: 575829. 10.3389/fpls.2020.575829

15. Garduño-Jiménez, A.-L., Carter, L.J., 2024. Insights into mode of action mediated responses following pharmaceutical uptake and accumulation in plants. Frontiers in Agronomy, 5: 1293555. 10.3389/fagro.2023.1293555

16. Gerlin, L., Cottret, L., Escourrou, A., Genin, S., Baroukh, C., 2022. A multi-organ metabolic model of tomato predicts plant responses to nutritional and genetic perturbations. Plant Physiology 188, 1709–1723. 10.1093/plphys/kiab548

17. Gorovits, R., Shteinberg, M., Mishra, R., Ari, J.B., Malchi, T., Chefetz, B., Anfoka, G., Czosnek, H., 2021. Interplay of stress responses to carbamazepine treatment, whitefly infestation and virus infection in tomato plants. Plant Stress 1, 100009. 10.1016/j.stress.2021.100009

18. Gorovits, R., Sobol, I., Akama, K., Chefetz, B., Czosnek, H., 2020. Pharmaceuticals in treated wastewater induce a stress response in tomato plants. Scientific Reports 10, 1856. 10.1038/s41598-020-58776-z

19. Groot, C.C. de, Marcelis, L.F.M., Boogaard, R. van den, Lambers, H., 2002. Interactive effects of nitrogen and irradiance on growth and partitioning of dry mass and nitrogen in young tomato plants. Functional Plant Biology 29, 1319–1328. 10.1071/fp02087

20. Guan, C., Cui, X., Liu, H., Li, X., Li, M., Zhang, Y., 2020. Proline Biosynthesis Enzyme Genes Confer Salt Tolerance to Switchgrass (*Panicum virgatum* L.) in Cooperation With Polyamines Metabolism. Frontiers in Plant Science 11:46. 10.3389/fpls.2020.00046

21. Hasan, M.M., Skalicky, M., Jahan, M.S., Hossain, M.N., Anwar, Z., Nie, Z.-F., Alabdallah, N.M., Brestic, M., Hejnak, V., Fang, X.-W., 2021. Spermine: Its Emerging Role in Regulating Drought Stress Responses in Plants. Cells 10, 261. 10.3390/cells10020261

22. Heirendt, L., Arreckx, S., Pfau, T., Mendoza, S.N., Richelle, A., Heinken, A., Haraldsdóttir, H.S., Wachowiak, J., Keating, S.M., Vlasov, V., Magnusdóttir, S., Ng, C.Y., Preciat, G., Žagare, A., Chan, S.H.J., Aurich, M.K., Clancy, C.M., Modamio, J., Sauls, J.T., Noronha, A., Bordbar, A., Cousins, B., El Assal, D.C., Valcarcel, L.V., Apaolaza, I., Ghaderi, S., Ahookhosh, M., Ben Guebila, M., Kostromins, A., Sompairac, N., Le, H.M., Ma, D., Sun, Y., Wang, L., Yurkovich, J.T., Oliveira, M.A.P., Vuong, P.T., El Assal, L.P., Kuperstein, I., Zinovyev, A., Hinton, H.S., Bryant, W.A., Aragón Artacho, F.J., Planes, F.J., Stalidzans, E., Maass, A., Vempala, S., Hucka, M., Saunders, M.A., Maranas, C.D., Lewis, N.E., Sauter, T., Palsson, B.Ø., Thiele, I., Fleming, R.M.T., 2019. Creation and analysis of biochemical constraint-based models using the COBRA Toolbox v.3.0. Nature protocols 14, 639–702. 10.1038/s41596-018-0098-2

23. Heberle, H., Meirelles, G.V., da Silva, F.R., Telles, G.P., Minghim, R., 2015. InteractiVenn: a web-based tool for the analysis of sets through Venn diagrams. BMC Bioinformatics 16, 169. 10.1186/s12859-015-0611-3

24. Herrmann, H.A., Dyson, B.C., Vass, L., Johnson, G.N., Schwartz, J.-M., 2019. Flux sampling is a powerful tool to study metabolism under changing environmental conditions. NPJ Systems Biology Applications 5, 32. 10.1038/s41540-019-0109-0

25. Hosseinifard, M., Stefaniak, S., Ghorbani Javid, M., Soltani, E., Wojtyla, Ł., Garnczarska, M., 2022. Contribution of Exogenous Proline to Abiotic Stresses Tolerance in Plants: A Review. International Journal of Molecular Sciences 23, 5186. 10.3390/ijms23095186

26. Jiménez-Arias, D., García-Machado, F.J., Morales-Sierra, S., García-García, A.L., Herrera, A.J., Valdés, F., Luis, J.C., Borges, A.A., 2021. A Beginner’s Guide to Osmoprotection by Biostimulants. Plants 10, 363. 10.3390/plants10020363

27. Kahlaoui, B., Hachicha, M., Rejeb, S., Rejeb, M.N., Hanchi, B., Misle, E., 2014. Response of two tomato cultivars to field-applied proline under irrigation with saline water: Growth, chlorophyll fluorescence and nutritional aspects. Photosynthetica 52, 421–429. 10.1007/s11099-014-0053-6

28. Kauffman, K.J., Prakash, P., Edwards, J.S., 2003. Advances in flux balance analysis. Current Opinion in Biotechnology 14, 491–496. 10.1016/j.copbio.2003.08.001

29. Kaya, C., Aydemir, S., Sonmez, O., Ashraf, M., Dikilitas, M., 2013. Regulation of growth and some key physiological processes in salt-stressed maize (*Zea mays* L.) plants by exogenous application of asparagine and glycerol. Acta Botanica Croatica 72, 157–168. 10.2478/v10184-012-0012-x

30. Knight, E.R., Carter, L.J., McLaughlin, M.J., 2018. Bioaccumulation, uptake, and toxicity of carbamazepine in soil-plant systems. Environmental Toxicology and Chemistry 37, 1122–1130. 10.1002/etc.4053

31. Knudsen, C., Gallage, N.J., Hansen, C.C., Møller, B.L., Laursen, T., 2018. Dynamic metabolic solutions to the sessile life style of plants. Natural Product Reports 35, 1140– 1155. 10.1039/c8np00037a

32. Lakshmanan, M., Cheung, C.Y.M., Mohanty, B., Lee, D.-Y., 2016. Modeling Rice Metabolism: From Elucidating Environmental Effects on Cellular Phenotype to Guiding Crop Improvement. Frontiers in Plant Science, 7:1795. 10.3389/fpls.2016.01795

33. Landa, P., Prerostova, S., Langhansova, L., Marsik, P., Vanek, T., 2017. Transcriptomic response of *Arabidopsis thaliana* (L.) Heynh. roots to ibuprofen. International Journal of Phytoremediation 19, 695–700. 10.1080/15226514.2016.1267697

34. Landa, P., Prerostova, S., Langhansova, L., Marsik, P., Vankova, R., Vanek, T., 2018. Transcriptomic response of *Arabidopsis thaliana* roots to naproxen and praziquantel. Ecotoxicology and Environmental Safety 166, 301–310. 10.1016/j.ecoenv.2018.09.081

35. Leitão, I., Leclercq, C.C., Ribeiro, D.M., Renaut, J., Almeida, A.M., Martins, L.L., Mourato, M.P., 2021a. Stress response of lettuce (*Lactuca sativa*) to environmental contamination with selected pharmaceuticals: A proteomic study. Journal of Proteomics 245, 104291. 10.1016/j.jprot.2021.104291

36. Leitão, I., Mourato, M. P., Carvalho, L., Oliveira, M. C., Marques, M. M., & Martins, L. L., 2021b. Antioxidative response of lettuce (*Lactuca sativa*) to carbamazepine-induced stress. Environmental Science and Pollution Research, 28, 45920–45932. 10.1007/s11356-021-13979-3

37. Li, Y., Qiu, L., Liu, X., Zhang, Q., Zhuansun, X., Fahima, T., Krugman, T., Sun, Q., Xie, C., 2020. Glycerol-Induced Powdery Mildew Resistance in Wheat by Regulating Plant Fatty Acid Metabolism, Plant Hormones Cross-Talk, and Pathogenesis-Related Genes. International Journal of Molecular Sciences 21, 673. 10.3390/ijms21020673

38. Li, Z., Cheng, B., Liu, W., Feng, G., Zhao, J., Zhang, L., Peng, Y., 2022. Global Metabolites Reprogramming Induced by Spermine Contributing to Salt Tolerance in Creeping Bentgrass. International Journal of Molecular Sciences 23, 4472. 10.3390/ijms23094472

39. Lieven, C., Beber, M.E., Olivier, B.G., Bergmann, F.T., Ataman, M., Babaei, P., Bartell, J.A., Blank, L.M., Chauhan, S., Correia, K., Diener, C., Dräger, A., Ebert, B.E., Edirisinghe, J.N., Faria, J.P., Feist, A.M., Fengos, G., Fleming, R.M.T., García-Jiménez, B., Hatzimanikatis, V., van Helvoirt, W., Henry, C.S., Hermjakob, H., Herrgård, M.J., Kaafarani, A., Kim, H.U., King, Z., Klamt, S., Klipp, E., Koehorst, J.J., König, M., Lakshmanan, M., Lee, D.-Y., Lee, S.Y., Lee, S., Lewis, N.E., Liu, F., Ma, H., Machado, D., Mahadevan, R., Maia, P., Mardinoglu, A., Medlock, G.L., Monk, J.M., Nielsen, J., Nielsen, L.K., Nogales, J., Nookaew, I., Palsson, B.O., Papin, J.A., Patil, K.R., Poolman, M., Price, N.D., Resendis-Antonio, O., Richelle, A., Rocha, I., Sánchez, B.J., Schaap, P.J., Malik Sheriff, R.S., Shoaie, S., Sonnenschein, N., Teusink, B., Vilaça, P., Vik, J.O., Wodke, J.A.H., Xavier, J.C., Yuan, Q., Zakhartsev, M., Zhang, C., 2020. MEMOTE for standardized genome-scale metabolic model testing. Nature Biotechnology 38, 272–276. 10.1038/s41587-020-0446-y

40. Lu, Z.-Y., Liu, C.-Y., Hu, Y.-Y., Pan, Y., Yuan, L., Wu, L.-T., Qi, K.-K., Zhang, Z., Zhou, J.-C., Zhao, J.-H., Hu, Y., Yin, H., Sheng, G.-P., 2024. Unmasking Spatial Heterogeneity in Phytotoxicology Mechanisms Induced by Carbamazepine by Mass Spectrometry Imaging and Multiomics Analyses. Environmental Science & Technology. 58, 13986–13994. 10.1021/acs.est.4c04628

41. M. Ali, R., Elfeky, S.S., Abbas, H., 2008. Response of Salt Stressed *Ricinus communis* L. To Exogenous Application of Glycerol and/or Aspartic Acid. Journal of Biological Sciences 8, 171–175. 10.3923/jbs.2008.171.175

42. Malchi, T., Eyal, S., Czosnek, H., Shenker, M., Chefetz, B., 2022. Plant pharmacology: Insights into in-planta kinetic and dynamic processes of xenobiotics. Critical Reviews in Environmental Science and Technology 52, 3525–3546. 10.1080/10643389.2021.1946360

43. Mascellani, A., Mercl, F., Kurhan, S., Pierdona, L., Kudrna, J., Zemanova, V., Hnilicka, F., Kloucek, P., Tlustos, P., Havlik, J., 2023. Biochemical and physiological changes in *Zea mays* L. after exposure to the environmental pharmaceutical pollutant carbamazepine. Chemosphere 329, 138689. 10.1016/j.chemosphere.2023.138689

44. Meech, R., Miners, J.O., Lewis, B.C., Mackenzie, P.I., 2012. The glycosidation of xenobiotics and endogenous compounds: Versatility and redundancy in the UDP glycosyltransferase superfamily. Pharmacology & Therapeutics 134, 200–218. 10.1016/j.pharmthera.2012.01.009

45. Nahar, K., Rahman, M., Hasanuzzaman, M., Alam, Md.M., Rahman, A., Suzuki, T., Fujita, M., 2016. Physiological and biochemical mechanisms of spermine-induced cadmium stress tolerance in mung bean (*Vigna radiata* L.) seedlings. Environmental Science and Pollution Research 23, 21206–21218. 10.1007/s11356-016-7295-8

46. Nanda, P., Patra, P., Das, M., Ghosh, A., 2020. Reconstruction and analysis of genome-scale metabolic model of weak Crabtree positive yeast *Lachancea kluyveri*. Scientific Reports 10, 16314. 10.1038/s41598-020-73253-3

47. Nguyen, H.M., Sako, K., Matsui, A., Suzuki, Y., Mostofa, M.G., Ha, C.V., Tanaka, M., Tran, L.-S.P., Habu, Y., Seki, M., 2017. Ethanol Enhances High-Salinity Stress Tolerance by Detoxifying Reactive Oxygen Species in *Arabidopsis thaliana* and Rice. Frontiers in Plant Science, 8:1001. 10.3389/fpls.2017.01001

48. Nguyen, M.-K., Lin, C., Nguyen, H.-L., Hung, N.T.Q., La, D.D., Nguyen, X.H., Chang, S.W., Chung, W.J., Nguyen, D.D., 2023. Occurrence, fate, and potential risk of pharmaceutical pollutants in agriculture: Challenges and environmentally friendly solutions. Science of The Total Environment 899, 165323. 10.1016/j.scitotenv.2023.165323

49. Ofaim, S., Zarecki, R., Porob, S., Gat, D., Lahav, T., Kashi, Y., Aly, R., Eizenberg, H., Ronen, Z., Freilich, S., 2020. Genome-scale reconstruction of *Paenarthrobacter aurescens* TC1 metabolic model towards the study of atrazine bioremediation. Scientific Reports 10, 13019. 10.1038/s41598-020-69509-7

50. Pascual, L.S., López-Climent, M.F., Segarra-Medina, C., Gómez-Cadenas, A., Zandalinas, S.I., 2023. Exogenous spermine alleviates the negative effects of combined salinity and paraquat in tomato plants by decreasing stress-induced oxidative damage. Frontiers in Plant Science, 14: 1193207. 10.3389/fpls.2023.1193207

51. Patel, M., Kumar, R., Kishor, K., Mlsna, T., Pittman, C.U.Jr., Mohan, D., 2019. Pharmaceuticals of Emerging Concern in Aquatic Systems: Chemistry, Occurrence, Effects, and Removal Methods. Chemical reviews 119, 3510–3673. 10.1021/acs.chemrev.8b00299

52. Poudel, S., Shrestha, A., Kandel, N., Adhikari, S., Paudel, S.R., 2023. A Review of Reclaimed Water Reuse for Irrigation in South Asian Countries. ACS EST Water 3, 3790–3806. 10.1021/acsestwater.3c00487

53. Rahman, M.M., Mostofa, M.G., Das, A.K., Anik, T.R., Keya, S.S., Ahsan, S.M., Khan, M.A.R., Ahmed, M., Rahman, M.A., Hossain, M.M., Tran, L.-S.P., 2022. Ethanol Positively Modulates Photosynthetic Traits, Antioxidant Defense and Osmoprotectant Levels to Enhance Drought Acclimatization in Soybean. Antioxidants 11, 516. 10.3390/antiox11030516

54. Raman, K., Chandra, N., 2009. Flux balance analysis of biological systems: applications and challenges. Briefings in Bioinformatics 10, 435–449. 10.1093/bib/bbp011

55. Raman, K., Kratochvíl, M., Olivier, B.G., König, M., Sengupta, P., Baskaran, D.K.K., Nguyen, T.V.N., Lobo, D., Wilken, S.E., Tiwari, K.K., Raghu, A.K., Palanikumar, I., Raajaraam, L., Ibrahim, M., Balakrishnan, S., Umale, S., Bergmann, F., Malpani, T., Satagopam, V.P., Schneider, R., Beber, M.E., Keating, S., Anton, M., Renz, A., Lakshmanan, M., Lee, D.-Y., Koduru, L., Mostolizadeh, R., Dias, O., Cunha, E., Oliveira, A., Lee, Y.Q., Zengler, K., Santibáñez-Palominos, R., Kumar, M., Barberis, M., Puniya, B.L., Helikar, T., Dinh, H.V., Suthers, P.F., Maranas, C.D., Casini, I., Loghmani, S.B., Veith, N., Leonidou, N., Li, F., Chen, Y., Nielsen, J., Lee, G., Lee, S.M., Kim, G.B., Monteiro, P.T., Teixeira, M.C., Kim, H.U., Lee, S.Y., Liebal, U.W., Blank, L.M., Lieven, C., Tarzi, C., Angione, C., Blaise, M.E., Aytar, Ç.P., Kulyashov, M., Akberdin, L., Kim, D., Yoon, S.H., Xu, Z., Gautam, J., Scott, W.T., Schaap, P.J., Koehorst, J.J., Zuñiga, C., Canto-Encalada, G., Benito-Vaquerizo, S., Olm, I.P., Suarez-Diez, M., Yuan, Q., Ma, H., Islam, M.M., Papin, J.A., Zorrilla, F., Patil, K.R., Basile, A., Nogales, J., León, G.S., Castillo-Alfonso, F., Olivares-Hernández, R., Canto-Encalada, G., Vigueras-Ramírez, G., Hermjakob, H., Dräger, A., Malik-Sheriff, R.S., 2024. FROG Analysis Ensures the Reproducibility of Genome Scale Metabolic Models. bioRxiv. 10.1101/2024.09.24.614797

56. Raoufi, A., Rahemi, M., Akbari, M., 2020. Glycerol foliar application improves salt tolerance in three pistachio rootstocks. Journal of the Saudi Society of Agricultural Sciences 19, 426–437. 10.1016/j.jssas.2020.07.003

57. Ravichandran, M.K., Philip, L., 2022. Fate of carbamazepine and its effect on physiological characteristics of wetland plant species in the hydroponic system. Science of The Total Environment 846, 157337. 10.1016/j.scitotenv.2022.157337

58. Raza, A., Charagh, S., Abbas, S., Hassan, M.U., Saeed, F., Haider, S., Sharif, R., Anand, A., Corpas, F.J., Jin, W., Varshney, R.K., 2023. Assessment of proline function in higher plants under extreme temperatures. Plant Biology 25, 379–395. 10.1111/plb.13510

59. Riemenschneider, C., Al-Raggad, M., Moeder, M., Seiwert, B., Salameh, E., Reemtsma, T., 2016. Pharmaceuticals, Their Metabolites, and Other Polar Pollutants in Field-Grown Vegetables Irrigated with Treated Municipal Wastewater. Journal of Agricultural and Food Chemistry 64, 5784–5792. 10.1021/acs.jafc.6b01696

60. Riemenschneider, C., Seiwert, B., Moeder, M., Schwarz, D., Reemtsma, T., 2017. Extensive Transformation of the Pharmaceutical Carbamazepine Following Uptake into Intact Tomato Plants. Environmental Science & Technology. 51, 6100–6109. 10.1021/acs.est.6b06485

61. Righetti, L., Rolli, E., Dellafiora, L., Galaverna, G., Suman, M., Bruni, R., Dall’Asta, C., 2021. Thinking Out of the Box: On the Ability of *Zea mays* L. to Biotrasform Aflatoxin B1 Into Its Modified Forms. Frontiers in Plant Science, 11: 599158. 10.3389/fpls.2020.c

62. Sahoo, S., Haraldsdóttir, H.S., Fleming, R.M.T., Thiele, I., 2015. Modeling the effects of commonly used drugs on human metabolism. The FEBS Journal 282, 297–317. 10.1111/febs.13128

63. Sandermann, H., Diesperger, H., Scheel, D., 1977. Metabolism of Xenobiotics by Plant Cell Cultures, in: Barz, W., Reinhard, E., Zenk, M.H. (Eds.), Plant Tissue Culture and Its Bio-Technological Application. Springer, Berlin, Heidelberg, pp. 178–196. 10.1007/978-3-642-66646-9_15

64. Scheel, D., Sandermann Jr., H., 1977. Metabolism of DDT and Kelthane in cell suspension cultures of parsley (*Petroselinum hortense*, Hoffm.) and soybean (Glycine max L.). Planta 133, 315–320. 10.1007/BF00380695

65. Schröder, P., 2007. Exploiting Plant Metabolism for the Phytoremediation of Organic Xenobiotics, in: Willey, N. (Ed.), Phytoremediation: Methods and Reviews. Humana Press, Totowa, NJ, pp. 251–263. 10.1007/978-1-59745-098-0_20

66. Shameer, S., Ratcliffe, R.G., Sweetlove, L.J., 2019. Leaf Energy Balance Requires Mitochondrial Respiration and Export of Chloroplast NADPH in the Light. Plant Physiology 180, 1947–1961. 10.1104/pp.19.00624

67. Shiade, S.R.G., Zand-Silakhoor, A., Fathi, A., Rahimi, R., Minkina, T., Rajput, V.D., Zulfiqar, U., Chaudhary, T., 2024. Plant metabolites and signaling pathways in response to biotic and abiotic stresses: Exploring bio stimulant applications. Plant Stress 12, 100454. 10.1016/j.stress.2024.100454

68. Thorn, C.F., Leckband, S.G., Kelsoe, J., Steven Leeder, J., Müller, D.J., Klein, T.E., Altman, R.B., 2011. PharmGKB summary: carbamazepine pathway. Pharmacogenetics and Genomics 21, 906. 10.1097/FPC.0b013e328348c6f2

69. Todaka, D., Quynh, D.T.N., Tanaka, M., Utsumi, Y., Utsumi, C., Ezoe, A., Takahashi, S., Ishida, J., Kusano, M., Kobayashi, M., Saito, K., Nagano, A.J., Nakano, Y., Mitsuda, N., Fujiwara, S., Seki, M., 2024. Application of ethanol alleviates heat damage to leaf growth and yield in tomato. Frontiers in Plant Science, 15: 1325365. 10.3389/fpls.2024.1325365

70. Van Oosten, M.J., Pepe, O., De Pascale, S., Silletti, S., Maggio, A., 2017. The role of biostimulants and bioeffectors as alleviators of abiotic stress in crop plants. Chemical and Biological Technologies in Agriculture 4, 5. 10.1186/s40538-017-0089-5

71. Varma, A., Palsson, B.O., 1994. Metabolic Flux Balancing: Basic Concepts, Scientific and Practical Use. Nature Biotechnology 12, 994–998. 10.1038/nbt1094-994

72. Voss, I., Sunil, B., Scheibe, R., Raghavendra, A.S., 2013. Emerging concept for the role of photorespiration as an important part of abiotic stress response. Plant Biology 15, 713–722. 10.1111/j.1438-8677.2012.00710.x

73. Wahman, R., Sauvêtre, A., Schröder, P., Moser, S., Letzel, T., 2020. Untargeted Metabolomics Studies on Drug-Incubated *Phragmites australis* Profiles. Metabolites 11, 2. 10.3390/metabo11010002

74. Wang, Y., Lu, J., Mao, L., Li, J., Yuan, Z., Bond, P.L., Guo, J., 2019. Antiepileptic drug carbamazepine promotes horizontal transfer of plasmid-borne multi-antibiotic resistance genes within and across bacterial genera. ISME J 13, 509–522. 10.1038/s41396-018-0275-x

75. Wanichthanarak, K., Boonchai, C., Kojonna, T., Chadchawan, S., Sangwongchai, W., Thitisaksakul, M., 2020. Deciphering rice metabolic flux reprograming under salinity stress via *in silico* metabolic modeling. Computational and Structural Biotechnology Journal 18, 3555–3566. 10.1016/j.csbj.2020.11.023

76. Wei, H., Tang, M., Xu, X., 2023. Mechanism of uptake, accumulation, transport, metabolism and phytotoxic effects of pharmaceuticals and personal care products within plants: A review. Science of the Total Environment 892, 164413. 10.1016/j.scitotenv.2023.164413

77. Whirl-Carrillo, M., Huddart, R., Gong, L., Sangkuhl, K., Thorn, C.F., Whaley, R., Klein, T.E., 2021. An Evidence-Based Framework for Evaluating Pharmacogenomics Knowledge for Personalized Medicine. Clinical Pharmacology & Therapeutics 110, 563–572. 10.1002/cpt.2350

78. Williams, T.C.R., Poolman, M.G., Howden, A.J.M., Schwarzlander, M., Fell, D.A., Ratcliffe, R.G., Sweetlove, L.J., 2010. A Genome-Scale Metabolic Model Accurately Predicts Fluxes in Central Carbon Metabolism under Stress Conditions. Plant Physiology 154, 311–323. 10.1104/pp.110.158535

79. Zhang, Y., Liu, J., Zhou, Y., Gong, T., Wang, J., Ge, Y., 2013. Enhanced phytoremediation of mixed heavy metal (mercury)–organic pollutants (trichloroethylene) with transgenic alfalfa co-expressing glutathione S-transferase and human P450 2E1. Journal of Hazardous Materials 260, 1100–1107. 10.1016/j.jhazmat.2013.06.065

