## Supplementary Text for "Investigating the impact of carbamazepine on tomato plant metabolism using genome-scale metabolic modelling"

### S1. Methods

#### S1.1 Curation of tomato GEM

The published model (MODEL2111120001) from the Biomodels database was converted to the MATLAB format by loading the SBML version of the model using COBRApy. The model reaction bounds were set to (-1000,1000) for reversible, and (0,1000) for irreversible reactions. The COBRA toolbox function 'checkMassChargeBalance' was used for evaluation of the mass and charge balance of the model. The stoichiometry and/or cofactor information for the reactions and the charge and/or formula for metabolites were updated using data from Chemical Entities of Biological Interest database (Hastings et al. 2016), Plant Metabolic Network databases (TomatoCyc, PlantCyc) (Hawkins et al. 2021), MetaCyc (Caspi et al. 2020) databases and prior published models (Shaw and Cheung 2019, 2021). A total of 1063 out of 1104 reaction mass and charge balance were fixed with the details of the unbalanced reactions provided in the supplementary excel file 'Supplementary_1.xlsx'. Further, to improve the robustness of the model, connectivity in the following pathways was fixed: sulphur metabolism, reactive oxygen species (ROS) generation, valine degradation, S-methyl-5'-thioadenosine degradation, pantothenate biosynthesis, isoleucine catabolism, leucine catabolism, lysine catabolism, and tyrosine metabolism.

To mimic the field conditions, phototrophic conditions as listed in Table S1 were constrained for further metabolic investigations. The non-growth associated maintenance (NGAM) constraint was fixed to 7.1 mmol $gDW^{-1}$ $h^{-1}$ as per the prior versions of tomato models (Gerlin et al. 2022; Yuan et al. 2016). For constraining the efficiency of photosynthesis and photorespiration, Ribulose-1,5-bisphosphate carboxylase/oxygenase (RubisCo) enzyme flux as carboxylase to oxygenase ratio was fixed as 85 as per prior version of tomato model (Gerlin et al. 2022). The upper bound of biomass reaction was constrained to an experimentally reported tomato plant-specific growth rate of 0.01 $h^{-1}$ (Gerlin et al. 2022; Groot et al. 2002).

**Table S1: Phototrophic uptake constraints**

| **Exchange reactions** | **Uptake constraints for maximum growth of leaf biomass as objective** |
|---|---|
| EX_mg2_b | (-1000,0) |
| EX_na_b | (-1000,0) |
| EX_nh4_b | (-1000,0) |
| EX_co2_b | (-1000,1000) |

| | |
|---|---|
| EX_h2o_b | (-1000,1000) |
| EX_pi_b | (-1000,0) |
| EX_so4_b | (-1000,0) |
| EX_cl_b | (-1000,0) |
| EX_k_b | (-1000,0) |
| EX_ca2_b | (-1000,0) |
| EX_no3_b | (-1000,0) |
| EX_photon_b | (-1000,0) |
| EX_o2_b | (-1000,1000) |
| ATPS (NGAM) | (7.1,1000) |
| Leaf biomass reaction | (0,0.01) |
| RubisCo flux as Carboxylase to Oxygenase ratio | $V_{RBPCh} / V_{RBCh\_1} = 85$ |

### S1.2 Curation of CBZ pharmacokinetics module

Among the emerging pharmaceutical pollutants, CBZ transformation has been extensively investigated in various plant species (Wei et al. 2023). Hence, the CBZ pharmacokinetic module was integrated into the updated GEM specific to the transformation metabolites reported in tomato plants (Riemenschneider et al. 2017). To represent these transformations in the model, the corresponding reactions of transformed metabolites were added to the model based on the 'green-liver' concept, as the reactions of CBZ transformation were not available in plant-specific biochemical databases. Cytochrome-P450 mediated transformation of phase-I metabolism was added based on animal liver metabolism of CBZ from Kyoto Encyclopedia of Genes and Genomes (KEGG) (Kanehisa et al. 2023) and Human Metabolome database (HMDB) (Wishart et al. 2022). Further, phase-II and phase-III metabolic conversions were manually curated in the model with metabolite formula adapted from the mass spectral data of CBZ transformation (Sauvêtre et al. 2018) with biochemical reaction adapted from TomatoCyc database (Hawkins et al. 2021). Transformations with unknown co-factor information in KEGG and HMDB were excluded in this study for a few phase-I experimentally reported CBZ metabolites in tomato such as 10,11-dihydrocarbamazepine and 10-hydroxycarbamazepine. Also, the acridine-mediated pathway of CBZ transformation has been ruled out for estimation of the metabolic burden of CBZ detoxification in tomato as it has been reported under strong oxidative physio-chemical conditions (Seiwert et al. 2015) and in abiotic control systems without plant counterparts (Sauvêtre et al. 2018; De Mastro et al. 2024). Based on experimental observations, CBZ, and its transformed metabolites have been detected in both in cell extract and in the spent media (Sauvêtre et al. 2018), and also to avoid making the transformed metabolites as dead-ends, the transformed metabolites were restricted to be secreted to extracellular space. Thus, 27 metabolites and 40 reactions (14 metabolic + 15 transport + 11 exchange reactions) were added (Figure S1) to the base tomato model to represent CBZ pharmacokinetics in tomato. Out of the 14 metabolic reactions, six reactions of phase I, five reactions of phase II, and three reactions of phase III metabolism of CBZ were added (Figure S1).

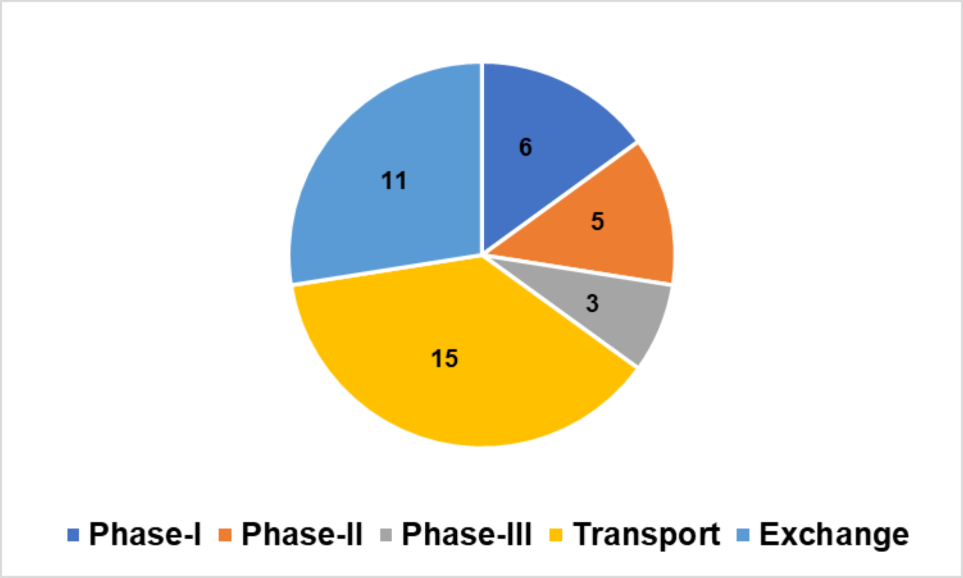


**Figure S1: Division of CBZ transformation reactions.**

#### S1.3 Algorithm for reaction classification

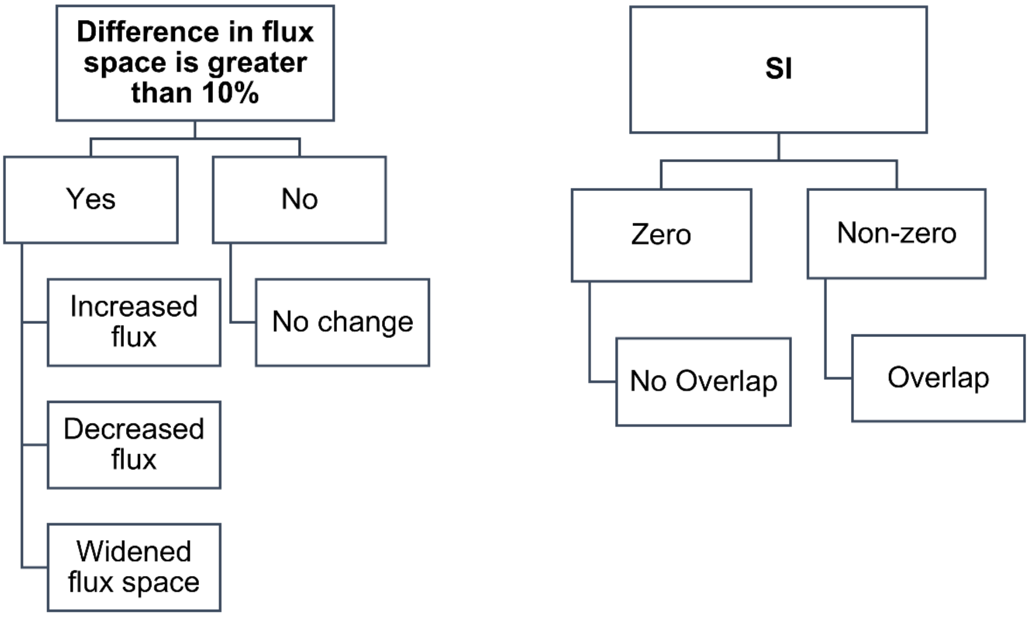


**Figure S2: Flow diagram representing the algorithm for classification of reactions into categories based on FVA results.**

### S1.4 Simulation of production capacity of metabolites

The maximum flux value from FVA for an exchange metabolite was used as an indicator of the production capability of the metabolite, and the minimum flux value from FVA was used as an indicator of the nutrient uptake capability in the model. Briefly, to simulate the production capacity of individual biomass components, the components of the leaf biomass equation were divided into categories of amino acids, nucleic acids, lipids, minerals, chlorophyll, sugar/organic acids, and lignin/(hemi)cellulose precursors as per Gerlin et al. (2022), and the average maximum flux under FVA for their corresponding exchange reaction (or minimum flux for input nutrient uptake-based exchange reactions) was calculated for a particular category. These values were compared for CBZ_IC50 condition in presence and absence of biostimulants.

### S1.5 Simulation for screening for key metabolites under CBZ stress response

Key metabolites with the potential to alleviate CBZ stress were screened using model simulations. Each of the individual metabolites represented in the biomass equation and a few major components of core metabolism such as Ribulose-5-bisphosphate, Erythrose-4-phosphate, Chorsimate, Ribose-5- phosphate, Pyruvate, Oxaloacetate, 3-phosphoglycerate, Dihydroxyacetone phosphate, Glyceraldehyde-3-phosphate, Fructose-1,6-bisphosphate, 6-phosphoglucanolactone and Glucose-6-phosphate were considered for the analysis. Under CBZ_IC50 condition with input constraints as per Table S1, for each of these metabolites, the lower bound of its exchange reaction was changed from 0 to -1000 mmol $gDW^{-1}$ $h^{-1}$ to simulate its exogenous addition. FBA (*l1*-norm) was estimated with maximizing biomass reaction as an objective. This analysis was performed iteratively for all these metabolites. A metabolite was considered to have the potential to alleviate CBZ stress if, under CBZ_IC50 condition, upon its exogenous addition, the specific growth rate was found to be increased with a threshold of at least 10 % improvement. Also, fold changes of key components of core metabolism screened in this study were compared for CBZ_IC50 condition in the presence and absence of biostimulants.

### S2. Evaluation of consistency of model prediction

#### S2.1 Evaluation of Consistency of updated model

Phototrophic conditions were simulated and the objective was fixed to obtain maximum phototrophic growth for leaf biomass. Table S2 summarises the results of the evaluation of the updated model in predicting the genetic perturbations, focusing on Sucrose phosphate synthase (SPS) gene knockout, attenuation of mitochondrial citrate synthase (MCS), attenuation of mitochondrial alpha-ketoglutarate dehydrogenase (AKGD), and attenuation of mitochondrial succinyl-CoA ligase (SUCOAS). The model accurately predicted the dependency on SPS flux for growth, while MCS, AKGD, and SUCOAS attenuation resulted in minor growth effects, consistent with unaltered leaf photosynthesis. However, discrepancies arose in the model's predictions of altered flux in nitrate metabolism for MCS attenuation. Also, GABA activation was observed in both control and mutant conditions for attenuation of AKGD and SUCOAS, not specific to the mutant. Further, the study investigated the reliability of the GEM module in predicting consistent behaviour in phototrophic modes. The updated GEM module consistently predicted activities in the phototrophic mode, with active ribulose-5-phosphate kinase and RubisCO, and zero as a possible solution for flux from α-ketoglutarate to malate under light conditions. The discrepancy between the predicted and experimental values can be reduced by fixing the blocked reactions /dead-end metabolites and further by constraining the model with experimental data. Further, with basic sanity checks from COBRA toolbox, the model was tested to be leakfree and doesn't produce energy with water and oxygen. Also, the model was tested to not produce matter when ATP demand reaction was reversed, ensuring biological consistency.

**Table S2 : Evaluation results of consistency of updated tomato model**

| S.No | Enzyme<br><br>Activity | Effect | CBZ _*i*SL3433 vs Experimental Observations |
|---|---|---|---|
| 1 | Sucrose phosphate synthase<br><br>Knockout<br><br>LB_mutant=UB_mutant =0<br><br>(Gerlin et al. 2022) | Growth Infeasible | Consistent |
| 2 | Mitochondrial citrate synthase<br><br>Attenuated<br><br>UB_mutant =<br><br>0.5 x UB_control<br><br>(Gerlin et al. 2022) | Enhanced CSp activity | Consistent |
| | | Minor effect on growth | Consistent |
| | | Reduced nitrate metabolism | Inconsistent (No Change) |
| | | Unaltered leaf photosynthesis | Consistent |
| 3 | Mitochondrial alpha-ketoglutarate dehydrogrenase<br><br>Attenuated | Unaltered leaf photosynthesis | Consistent |
| | | Minor effect on growth | Consistent |

| | | | |
|---|---|---|---|
| | UB_mutant = 0.5 x UB_control (Gerlin et al. 2022) | GABA shunt use | Inconsistent (Active in control and mutant conditions) |
| 4 | Mitochondrial succinyl-CoA ligase Attenuated UB_mutant = 0.5 x UB_control (Gerlin et al. 2022) | Unaltered leaf photosynthesis | Consistent |
| | | Minor effect on growth | Consistent |
| | | GABA shunt use | Inconsistent (Active in control and mutant conditions) |
| 5 | Phototrophic mode (Yuan et al. 2016) | Ribulose-5-phosphate kinase active | Consistent |
| | | RubisCO active | Consistent |
| | | α-ketoglutarate to malate carry no flux under light conditions | Consistent (Zero is there as solution) |

**S2.2 Prediction and verification of impact of nitrogen limitation on specific growth rate**

Under phototrophic conditions as per Table S1, the FBA (*l1*-norm) flux value of nitrate uptake flux under control conditions for maximizing biomass objective was considered as 100 %

optimum. With nitrate as the sole nitrogen source under phototrophic control conditions (with zero input of CBZ uptake rate) as per Table S1, the nitrate uptake rate was varied from 10 % to 100 % of optimum. For this input, FBA (*l1*-norm) with maximizing biomass reaction as objective function was estimated. The experimental values for nitrogen limitation on biomass growth rate were referenced from Groot et al. (2002). The values for the Virtual Young TOmato Plant (VYTOP) prediction were referenced from Gerlin et al. (2022). The results of CBZ_*i*SL3433, experimental values, and VYTOP predictions were compared, as represented in Figure S3.

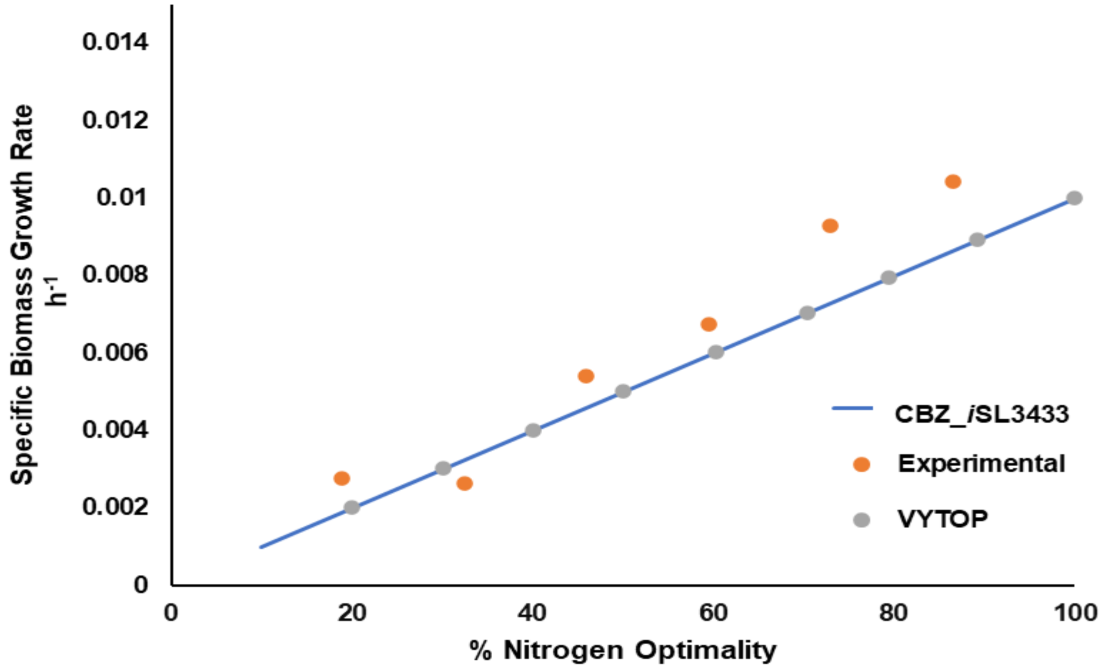


**Figure S3: Simulation of impact of nitrogen limitation on specific growth rate of leaf biomass. Growth rates were predicted with a limited nitrogen uptake flux using CBZ _*i*SL3433. This was compared with VYTOP model predictions and experimental data from Groot et al. (2002).**

### S3 Verification of consistency of prediction of significantly altered reaction set

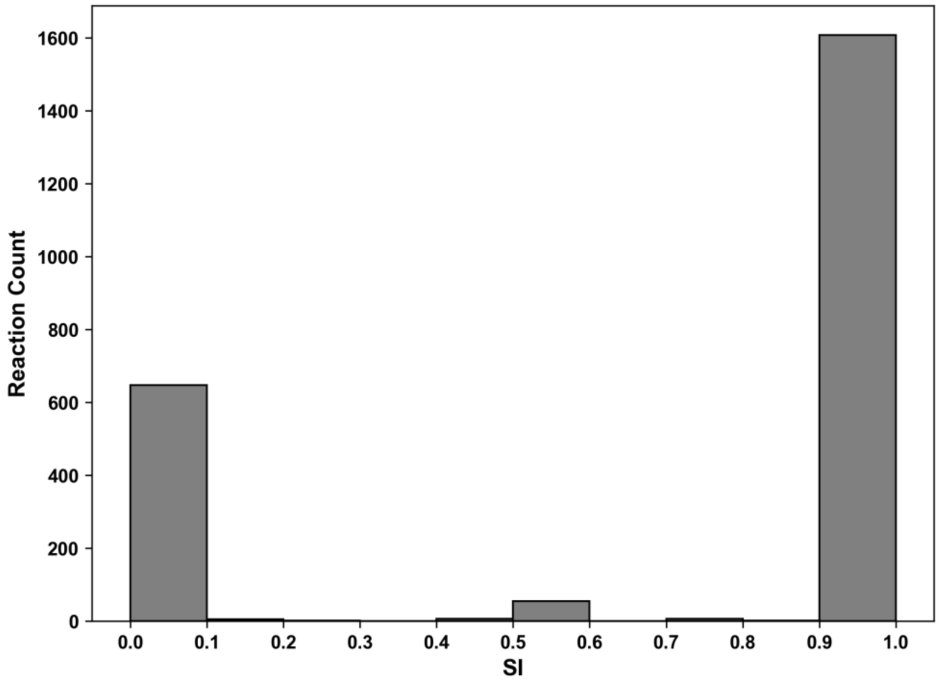


**Figure S4: SI distribution of flux spaces of reactions in CBZ_*i*SL3433 compared between Control and CBZ_IC50 conditions with minimizing photon uptake as objective.**

With biomass production as the objective, the FVA simulation was repeated for different CBZ uptake rates CBZ_IC25 and CBZ_IC75, representing uptake flux of CBZ inducing 25 % and 75 % reduction in specific growth rate compared to control conditions respectively. As observed from Figure S5A, 154 reactions estimated with CBZ_IC50 were a subset of CBZ_IC25 and CBZ_IC75, confirming significant alterations predicted for these 154 reactions were irrespective of CBZ uptake rates. Further, for verifying the independency of predicted alterations based on objective function choice, the FVA simulation was repeated with minimizing photon uptake as the objective function. Along with constraints as per Table S1, for control conditions, the biomass value was pinned to 0.01 $h^{-1}$, and for CBZ_IC50, the biomass value was pinned to 0.005 $h^{-1}$. FVA was simulated with minimizing photon uptake as objective and SI was estimated for all reactions. Approximately 27.79 % (647/2328) reactions had their SI in the range of 0 to 0.1 (Figure S4). Among these, 285 reactions were identified as significantly altered reactions with zero SI, and 99 % of 154 reactions previously identified as significantly altered with maximum biomass as objective function, were a part of this, thus signifying the predicted altered reactions were independent of the objective function choice of maximizing specific growth rate and minimizing photon uptake (Figure S5B).

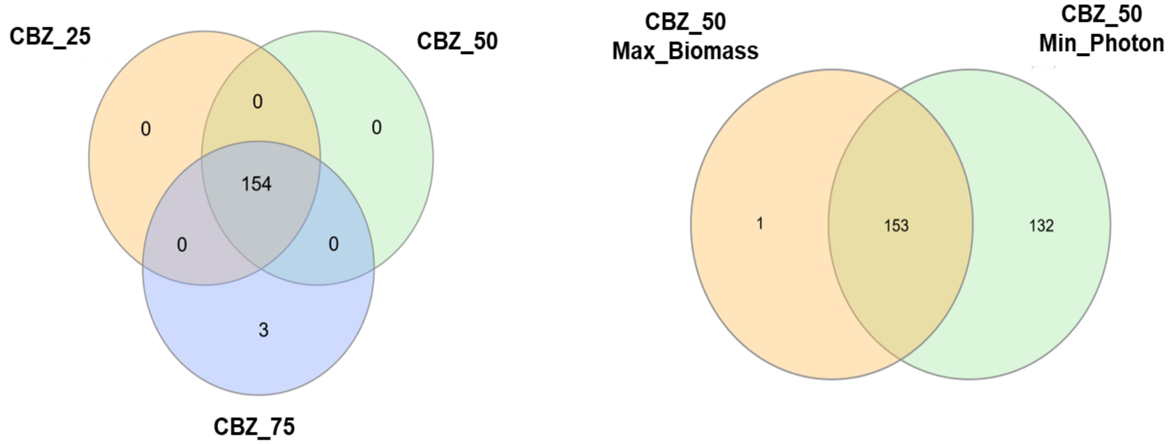


A B

**Figure S5: Venn diagram (Heberle et al. 2015) representing that the prediction of 154 significant altered reactions is independent of CBZ uptake rates (A), Objective function choice (B). A). Representation of intersection of sets of significantly altered reactions with increasing CBZ uptake rates. CBZ_25, CBZ_50, CBZ_75 each represents zero SI reaction sets of FVA between Control conditions and CBZ inducing 25 %, 50 %, 75 % reduction of biomass from control conditions. B). Representation of intersection of sets of significantly altered reactions predicted based on varied objective function optimization. Zero SI reaction sets of FVA between control and CBZ_IC50 conditions with 'CBZ_50 Max_Biomass' representing maximizing biomass production as objective function and with 'CBZ_50 Min_Photon' representing minimizing photon uptake as objective function.**

**S4 Comparison of transcriptomics data vs CBZ_*i*SL3433 prediction**

**Table S3: Comparison of q-PCR observation (Lu et al. 2024) vs CBZ_*i*SL3433 prediction for tomato plant exposed to CBZ**

| Gene identifier in transcriptome | Gene identifier in model | Reaction ID in the model | Enzyme Identity | SI | Diff>10 % in flux space | Predicted vs Experimental |
|---|---|---|---|---|---|---|
| **PHOTOSYNTHESIS** | | | | | | |
| SOLYC03G115980.1 | SOLYC03G115980.1 | DHGGDPR | 1.3.1.- | $6.99 \times 10^{-5}$ | Yes | Yes |
| | | DHGRGRCHLAR | | $6.99 \times 10^{-5}$ | Yes | Yes |
| | | GGDPR2h | | $6.99 \times 10^{-5}$ | Yes | Yes |
| | | GRGRCHLAR | | $6.99 \times 10^{-5}$ | Yes | Yes |
| SOLYC12G099930.2 | SOLYC12G099930.1 | AGTix | Alanine--glyoxylate aminotransferase | $8.33 \times 10^{-6}$ | Yes | Yes |
| | | GLYTAx | Glycine aminotransferase | $8.33 \times 10^{-6}$ | Yes | Yes |

| | | | | | | |
|---|---|---|---|---|---|---|
| | | SGATx | Serine--glyoxylate aminotransferase | $8.33 \times 10^{-6}$ | Yes | Yes |
| SOLYC01G097460.3 | SOLYC01G097460.2 | RPI | Ribose 5-phosphate epimerase | $7.35 \times 10^{-6}$ | Yes | Yes |
| | | RPIh | Ribose 5-phosphate epimerase | $6.21 \times 10^{-6}$ | Yes | Yes |
| SOLYC10G054870.2 | SOLYC10G054870.1 | TPI | Triosephosphate isomerase | 1 | No | No |
| | | TPIh | Triosephosphate isomerase | 1 | No | No |
| SOLYC12G094640.2 | SOLYC12G094640.1 | GAPD | Glyceraldehyde 3-phosphate dehydrogenase (phosphorylating) | 1 | No | No |
| | | GAPDH_nadp_hi | Glyceraldehyde-3-phosphate dehydrogenase (NADP+) (phosphorylating) | 0.56 | Yes | Yes |

| | | | | | | |
|---|---|---|---|---|---|---|
| | | GAPDHh | Glyceraldehyde 3-phosphate dehydrogenase (phosphorylating) | 1 | No | No |
| **ANTIOXIDANT EXPRESSION** | | | | | | |
| SOLYC12G094620.2 | SOLYC12G094620.1 | CAT2 | Catalase | $4.46 \times 10^{-7}$ | Yes | No |
| | | CATp | Catalase | $4.63 \times 10^{-6}$ | Yes | No |
| SOLYC01G067740.3 | SOLYC01G067740.2 | SOD_h | Superoxide dismutase | $2.56 \times 10^{-7}$ | Yes | No |
| | | SPODM | Superoxide dismutase | $2.56 \times 10^{-7}$ | Yes | No |

**Table S4: Comparison of transcriptomics (Lu et al. 2024) vs CBZ_*i*SL3433 prediction**

| S.No | Pathway | Gene identifier | Enzyme | Reaction ID in model | SI |
|---|---|---|---|---|---|
| 1 | Starch and Sucrose Metabolism | SOLYC10G046840.1 | Glycogen synthase | STARCH300S2 | 0.31 |
| 2 | Glutathione Metabolism | SOLYC09G065900.2 | Glutathione Reductase (Cytosol) | GTHOr | $2.56 \times 10^{-7}$ |
| | | | Glutathione Reductase (Mitochondria) | GTHOm | $2.56 \times 10^{-7}$ |
| | | | Glutathione Reductase (Peroxisome) | GTHOx | $4.61 \times 10^{-7}$ |
| | | | Glutathione Reductase (Plastid) | GDR_nadp_h | $3.97 \times 10^{-6}$ |

### S5 Altered Metabolism with CBZ stress

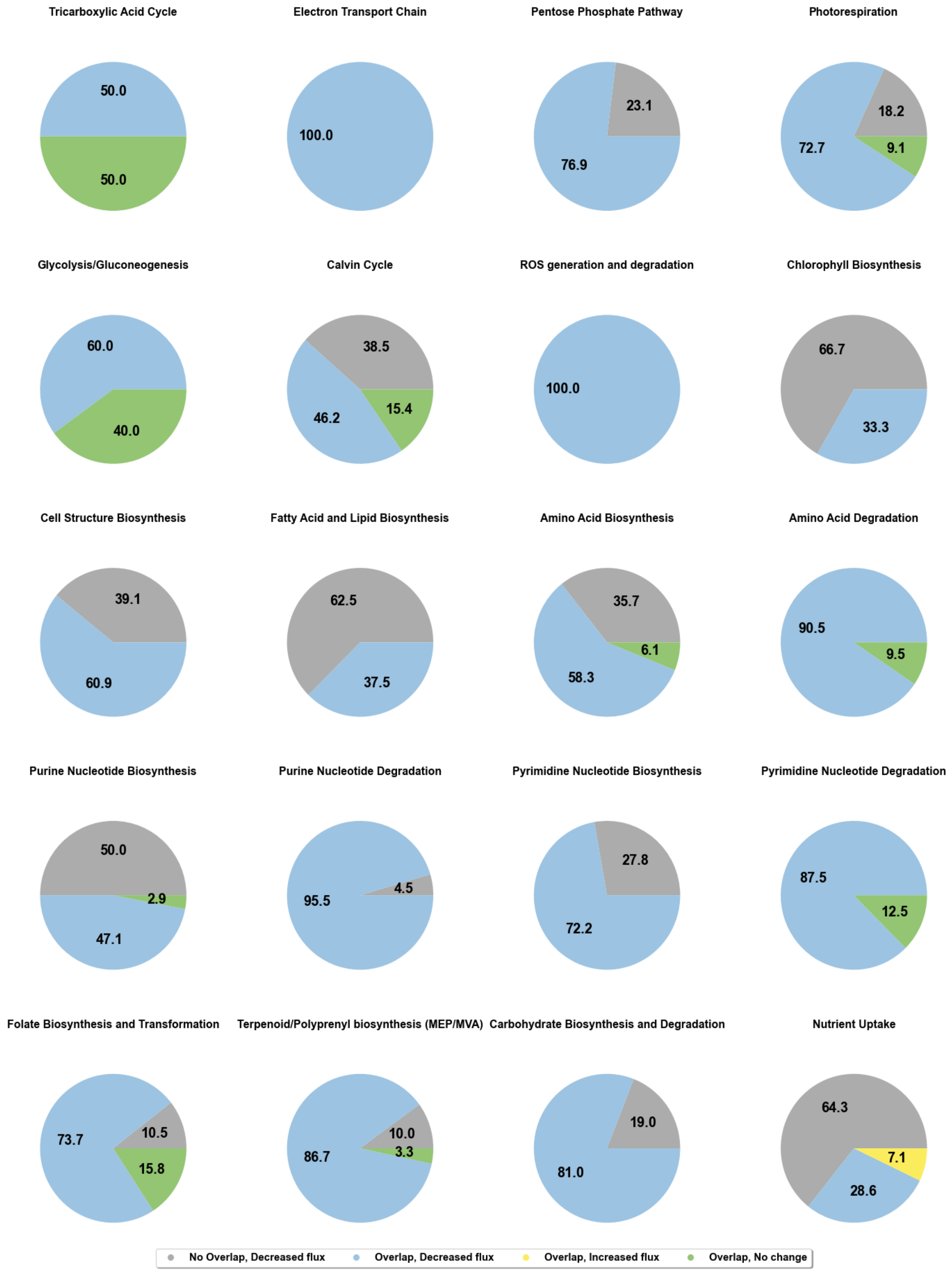

**Figure S6: Each pie-chart represents reprogramming of a specific metabolic pathway class in presence of CBZ. Numbers represent percentage of altered reactions in a pathway class. Each component of pie-chart represents one of the categories as mentioned in legend. Each category represents the classification of a reaction based on FVA analysis between Control and CBZ_IC50 condition as mentioned in Section 2.3.2.**

**S6 Effect of Biostimulants on ameliorative effect for CBZ stress**

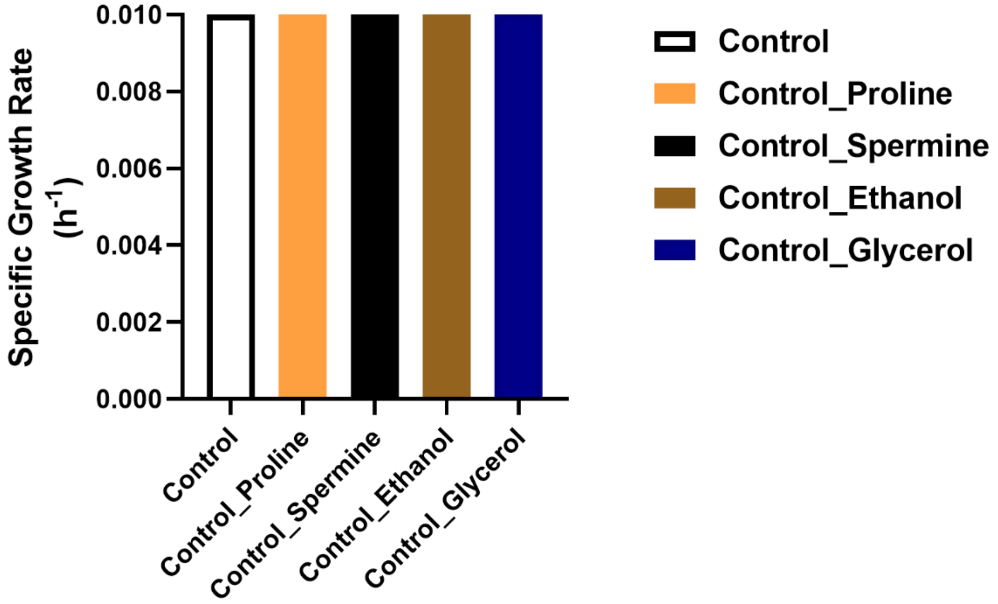


**Figure S7: Effect of exogenous addition of 0.07 mmol gDW$^{-1}$ h$^{-1}$ of biostimulants (Proline, Spermine, Ethanol and Glycerol) on specific growth rate under Control condition.**

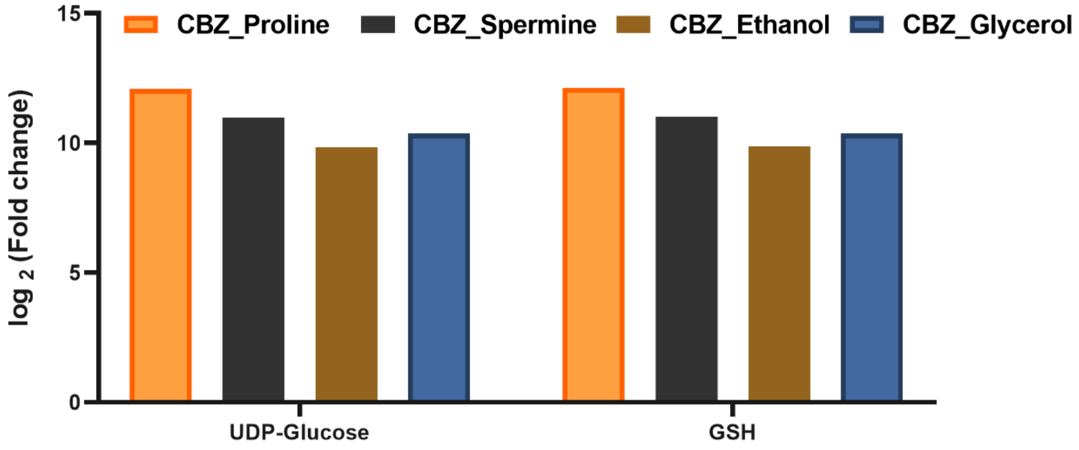


**Figure S8: Fold change ((CBZ_IC50_Biostimulant) / (CBZ_IC50)) of production capacity of UDP-Glucose and Glutathione with exogenous addition of 0.07 mmol $gDW^{-1}$ $h^{-1}$ of biostimulants. Fold-change values were transformed into logarithm base 2.**

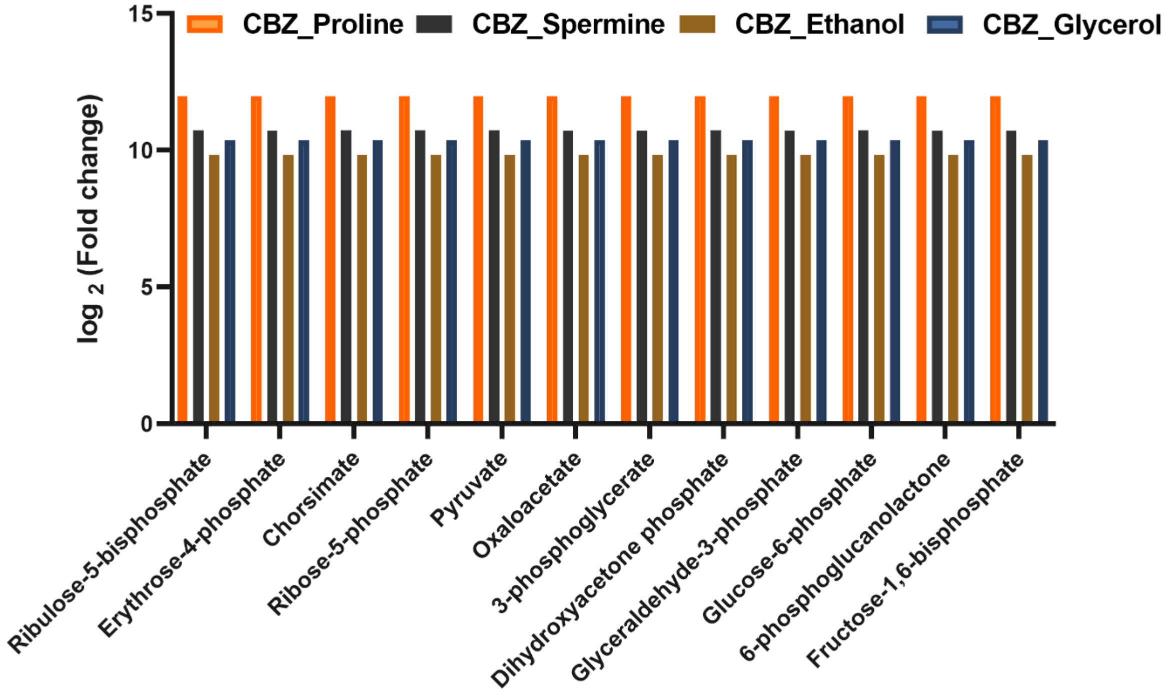


**Figure S9: Fold change ((CBZ_IC50_Biostimulant) / (CBZ_IC50)) of production capacity of key metabolites of core pathways with exogenous addition of 0.07 mmol $gDW^{-1}$ $h^{-1}$ of biostimulants. Fold-change values were transformed into logarithm base 2.**

**References:**


1. Caspi, R., Billington, R., Keseler, I.M., Kothari, A., Krummenacker, M., Midford, P.E., Ong, W.K., Paley, S., Subhraveti, P., Karp, P.D., 2020. The MetaCyc database of metabolic pathways and enzymes - a 2019 update. Nucleic Acids Research 48, D445–D453. https://doi.org/10.1093/nar/gkz862
2. De Mastro, F., Traversa, A., Cocozza, C., Cacace, C., Provenzano, M.R., Vona, D., Sannino, F., Brunetti, G., 2024. Fate of Carbamazepine and Its Metabolites in a Soil–Aromatic Plant System. Soil Systems 8, 83. https://doi.org/10.3390/soilsystems8030083
3. Gerlin, L., Cottret, L., Escourrou, A., Genin, S., Baroukh, C., 2022. A multi-organ metabolic model of tomato predicts plant responses to nutritional and genetic perturbations. Plant Physiology 188, 1709–1723. https://doi.org/10.1093/plphys/kiab548
4. Groot, C.C. de, Marcelis, L.F.M., Boogaard, R. van den, Lambers, H., 2002. Interactive effects of nitrogen and irradiance on growth and partitioning of dry mass and nitrogen in young tomato plants. Functional Plant Biology 29, 1319–1328. https://doi.org/10.1071/fp02087
5. Hastings, J., Owen, G., Dekker, A., Ennis, M., Kale, N., Muthukrishnan, V., Turner, S., Swainston, N., Mendes, P., Steinbeck, C., 2016. ChEBI in 2016: Improved services and an expanding collection of metabolites. Nucleic Acids Research 44, D1214-1219. https://doi.org/10.1093/nar/gkv1031
6. Hawkins, C., Ginzburg, D., Zhao, K., Dwyer, W., Xue, B., Xu, A., Rice, S., Cole, B., Paley, S., Karp, P., Rhee, S.Y., 2021. Plant Metabolic Network 15: A resource of genome-wide metabolism databases for 126 plants and algae. Journal of Integrative Plant Biology 63, 1888–1905. https://doi.org/10.1111/jipb.13163

7. Heberle, H., Meirelles, G.V., da Silva, F.R., Telles, G.P., Minghim, R., 2015. InteractiVenn: a web-based tool for the analysis of sets through Venn diagrams. BMC Bioinformatics 16, 169. https://doi.org/10.1186/s12859-015-0611-3
8. Kanehisa, M., Furumichi, M., Sato, Y., Kawashima, M., Ishiguro-Watanabe, M., 2023. KEGG for taxonomy-based analysis of pathways and genomes. Nucleic Acids Research 51, D587–D592. https://doi.org/10.1093/nar/gkac963
9. Lu, Z.-Y., Liu, C.-Y., Hu, Y.-Y., Pan, Y., Yuan, L., Wu, L.-T., Qi, K.-K., Zhang, Z., Zhou, J.-C., Zhao, J.-H., Hu, Y., Yin, H., Sheng, G.-P., 2024. Unmasking Spatial Heterogeneity in Phytotoxicology Mechanisms Induced by Carbamazepine by Mass Spectrometry Imaging and Multiomics Analyses. Environmental Science & Technology. 58, 13986–13994. https://doi.org/10.1021/acs.est.4c04628
10. Riemenschneider, C., Seiwert, B., Moeder, M., Schwarz, D., Reemtsma, T., 2017. Extensive Transformation of the Pharmaceutical Carbamazepine Following Uptake into Intact Tomato Plants. Environmental Science & Technology. 51, 6100–6109. https://doi.org/10.1021/acs.est.6b06485
11. Sauvêtre, A., May, R., Harpaintner, R., Poschenrieder, C., Schröder, P., 2018. Metabolism of carbamazepine in plant roots and endophytic rhizobacteria isolated from *Phragmites australis*. Journal of Hazardous Materials 342, 85–95. https://doi.org/10.1016/j.jhazmat.2017.08.006
12. Seiwert, B., Golan-Rozen, N., Weidauer, C., Riemenschneider, C., Chefetz, B., Hadar, Y., Reemtsma, T., 2015. Electrochemistry Combined with LC-HRMS: Elucidating Transformation Products of the Recalcitrant Pharmaceutical Compound Carbamazepine Generated by the White-Rot Fungus *Pleurotus ostreatus*. Environmental Science & Technology 49, 12342–12350. https://doi.org/10.1021/acs.est.5b02229

13. Shaw, R., Cheung, C.Y.M., 2021. Integration of crop growth model and constraint-based metabolic model predicts metabolic changes over rice plant development under water-limited stress. *in silico* Plants 3, diab020. https://doi.org/10.1093/insilicoplants/diab020
14. Shaw, R., Cheung, C.Y.M., 2019. A mass and charge balanced metabolic model of *Setaria viridis* revealed mechanisms of proton balancing in C4 plants. BMC Bioinformatics 20, 357. https://doi.org/10.1186/s12859-019-2941-z
15. Wei, H., Tang, M., Xu, X., 2023. Mechanism of uptake, accumulation, transport, metabolism and phytotoxic effects of pharmaceuticals and personal care products within plants: A review. Science of the Total Environment 892, 164413. https://doi.org/10.1016/j.scitotenv.2023.164413
16. Wishart, D.S., Guo, A., Oler, E., Wang, F., Anjum, A., Peters, H., Dizon, R., Sayeeda, Z., Tian, S., Lee, B.L., Berjanskii, M., Mah, R., Yamamoto, M., Jovel, J., Torres-Calzada, C., Hiebert-Giesbrecht, M., Lui, V.W., Varshavi, Dorna, Varshavi, Dorsa, Allen, D., Arndt, D., Khetarpal, N., Sivakumaran, A., Harford, K., Sanford, S., Yee, K., Cao, X., Budinski, Z., Liigand, J., Zhang, L., Zheng, J., Mandal, R., Karu, N., Dambrova, M., Schiöth, H.B., Greiner, R., Gautam, V., 2022. HMDB 5.0: the Human Metabolome Database for 2022. Nucleic Acids Research 50, D622–D631. https://doi.org/10.1093/nar/gkab1062
17. Yuan, H., Cheung, C.Y.M., Poolman, M.G., Hilbers, P.A.J., van Riel, N.A.W., 2016. A genome-scale metabolic network reconstruction of tomato (*Solanum lycopersicum* L.) and its application to photorespiratory metabolism. The Plant Journal 85, 289–304. https://doi.org/10.1111/tpj.13075
